# Solution to what? Global assessment of nature-based solutions, urban challenges, and outcomes

**DOI:** 10.1101/2023.12.07.570577

**Authors:** Meng Li, Roy P. Remme, Peter M. Van Bodegom, Alexander P.E. Van Oudenhoven

## Abstract

In response to multiple societal challenges faced in cities, nature-based solutions (NbS) are gaining prominence as means to support sustainable and resilient urban planning. However, NbS are being implemented in cities around the globe without comprehensive evidence on their effectiveness in addressing urban challenges. Based on a systematic mapping methodology, we synthesized 547 empirical cases of NbS in 197 cities globally, yielding 799 outcomes encompassing biodiversity, health well-being, and regulating ecosystem services. To structure this evidence we developed an urban NbS classification and categories of urban challenges and outcomes. Effectiveness of NbS was assessed through synthesizing which urban challenges are addressed by NbS, which outcomes are generated, and how these outcomes perform compared to alternative solutions. Our analysis suggests that specific urban challenges were mostly linked to closely related outcomes, but rarely to multiple outcomes. Specifically, forests & trees and general parks were commonly used to enhance health and well-being, while grassland and gardens were applied to mitigate biodiversity loss. Furthermore, urban NbS generally yielded positive effects compared to non-NbS, particularly in relation to microclimate mitigation and mental health outcomes. However, we note a scarcity of evidence on multifunctional NbS, especially on studies that report multiple outcomes related to biodiversity and well-being simultaneously. Our study provides a foundation for further understanding NbS effectiveness and can inform urban planners and policymakers with measurable evidenced-based targets for the application of NbS.

## 1 Introduction

Cities worldwide are confronted with increasing environmental, social, and economic challenges that together threaten the resilience and sustainability of urban areas (Bush & Doyon, 2019; Keivani, 2010). These challenges include biodiversity loss, flooding after intense rainfall, the urban heat island effect, air pollution, public health issues, and rising social-economic inequality (Brink et al., 2016; Frantzeskaki et al., 2019; Oke et al., 2021). Increasing urbanization, interacting with climate change, likely amplifies the existing challenges and generates new ones, leading to greater economic, social, and environmental losses in cities (Bazrkar et al., 2015). Given these growing threats to urban populations, nature-based solutions (NbS) have been suggested to provide integrated and multifunctional solutions to enhance resilience and sustainability in cities (Nesshöver et al., 2017).

Although definitions vary, NbS generally refer to actions that are inspired or powered by nature to address sustainability challenges and benefit both people and nature (Cohen-Shacham et al., 2016; European Commission, 2015). A key characteristic of NbS is their capacity to simultaneously benefit human well-being and biodiversity (Kabisch et al., 2022; Seddon et al., 2021; Sowińska-Świerkosz & García, 2022). Moreover, NbS target real-world issues explicitly with a problem– and objective-oriented approach (Dorst et al., 2019). In urban contexts, NbS approaches facilitate the integration of related concepts such as urban ecosystem services, green infrastructure and ecosystem-based approaches, to address urban sustainability challenges (henceforth urban challenges) (Fang et al., 2023; Remme et al., 2024). Furthermore, the focus of NbS on concrete actions and applications is seen as a core strength to connect urban challenges to policy and planning (Albert et al., 2019; Coletta et al., 2021; Wang et al., 2021). As a result, NbS have become more prominent in urban planning, policy, and research (Seddon et al., 2021).

The widespread adoption of NbS is hindered by uncertainty around their effectiveness to address urban challenges (Seddon, 2022). This uncertainty hampers the uptake of NbS by decision-makers and is one of the key challenges in mainstreaming NbS in cities (Frantzeskaki et al., 2019; Sarabi et al., 2019). This challenge is due to a lack of evidence base on NbS effectiveness across widespread contexts (Sarabi et al., 2019; Seddon et al., 2020). Insufficient evidence on effectiveness could lead to misunderstanding and misuse of NbS (Krauze & Wagner, 2019), such as the over-reliance on tree planting rather than a wide range of NbS, or simply ‘greening’ the city rather than considering the associated challenges and potential outcomes (Escobedo et al., 2018; Holl & Brancalion, 2020). Successful uptake and mainstreaming of NbS requires comprehensive evidence on the effectiveness of NbS to address urban challenges and provide multiple benefits.

Overall, key elements for assessing NbS effectiveness in cities include the urban challenges addressed, NbS applied, and actual outcomes assessed. Here, we understand NbS effectiveness in cities as the extent to which NbS address urban challenges and provide outcomes for biodiversity and human well-being. So far, the debate has mostly focused on the types of NbS, i.e. the implemented actions to address targeted challenges, and to what extent outcomes are achieved (Nature editorial, 2017; Raymond, Berry, et al., 2017; Seddon et al., 2020). However, to truly assess effectiveness, insights in the extent to which outcomes are provided is required, particularly compared to alternative solutions (Frantzeskaki et al., 2019; Sowińska-Świerkosz & García, 2021). Such assessments of outcomes should explicitly consider if multifunctionality can be achieved, benefitting both humans and biodiversity (Kabisch et al., 2022). This requires integrated assessments that capture multiple outcomes simultaneously (Key et al., 2022; Veerkamp et al., 2021). A scientific evidence base of NbS effectiveness, assessing these elements in combination, is essential for informing decision-making and management of NbS.

Notable contributions to this evidence base have been made in recent years, especially in the context of how various challenges are addressed by NbS. For example, Dunlop et al. (2024) found that NbS research has primarily focused on climate change and biodiversity loss, with a nascent focus on other societal challenges since 2015. Furthermore, research on the linkages between ecosystem services (ES), specific NbS, and urban challenges is developing (Babí Almenar et al., 2021; Fang et al., 2024). These studies contribute to our understanding on how NbS in general can tackle urban challenges. To a lesser extent, they address how specific NbS can, through the delivery of ES, address urban challenges. Hence, a knowledge gap remains on how effective specific NbS types are in addressing urban challenges (Babí Almenar et al., 2021; Fang et al., 2024).

In addition to synthesizing various challenges, reviews on NbS effectiveness have also evaluated outcomes. Previous studies evaluated NbS as general ecosystem types or interventions in natural environments, with a predominant focus on outcomes related to climate change and water-related risks specific to certain geographical regions (Cheng et al., 2023; Villamayor-Tomas et al., 2024; Xu et al., 2023). For example, Acreman et al. (2021) found that restoration of forests and floodplain wetlands can reduce flood risk in the African context. The only global assessment of NbS outcomes to date involved interventions outside urban areas and focused on climate change adaptation (Chausson et al., 2020). When focusing on NbS in urban contexts, reviews have focused on outcomes related to specific sets of NbS types, often related to climate change or health issues (Adewuyi et al., 2023; Harvey et al., 2024; Obeng et al., 2023). For example, Esraz-Ul-Zannat et al. (2024) investigated the effectiveness of small-scale NbS in reducing flood risks in comparison to grey infrastructure. Furthermore, while ecological aspects of urban biodiversity are increasingly known, biodiversity outcomes are not adequately addressed in current NbS reviews and lack connection to urban planning (Knapp & MacIvor, 2023). Despite the growing evidence on NbS outcomes, knowledge remains scattered across challenges and outcome categories (Johnson et al., 2022). An integrative assessment of NbS outcomes has not been undertaken, and is particularly lacking on a global level.

To address these knowledge gaps, we used a systematic mapping methodology to consolidate the scientific evidence on the effectiveness of real-world NbS cases in cities worldwide. Drawing from this evidence, we aimed to systematically analyze the effectiveness of NbS to address urban challenges and provide outcomes for biodiversity and human well-being. To achieve this, we 1) studied relations between urban challenges, NbS types, and their outcomes, 2) evaluated the performance of NbS compared to non-NbS, and 3) assessed cases reporting multiple outcomes and their interrelationships. To facilitate such analysis, we also developed a classification of urban NbS, to allow comparison among studies and facilitate the transfer of information to support urban planning and policy. Through this research, we take important initial steps and provide fundamental knowledge toward better understanding the effectiveness of urban NbS. Our findings contribute to informing urban planning and policy, by offering a consistent NbS classification and insights into which urban challenges can be addressed by NbS, and what kind of corresponding outcomes can be expected.

## 2. Methods

This research was guided by the systematic mapping standards set by Collaboration for Environmental Evidence (2018). It includes five phases: (1) defining the scope of the research, (2) conducting a systematic search to identify relevant studies, (3) screening for eligibility, (4) coding and extracting relevant data for NbS case studies identified and finally (5) analyzing the data (Fig. 1).

**Fig. 1.**
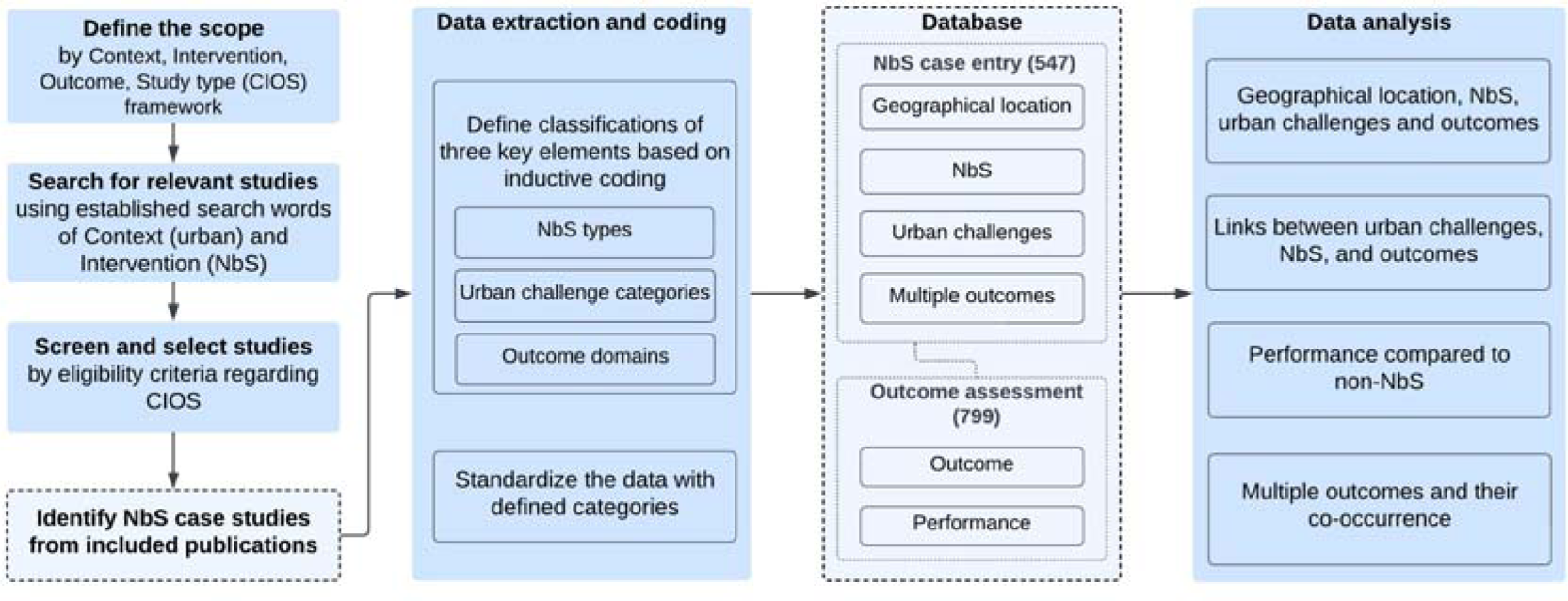
Schematic overview of the systematic mapping methodology. Methodological steps are depicted in borderless boxes, while data outputs are represented by dashed line-bordered boxes.

**Fig. 2.**
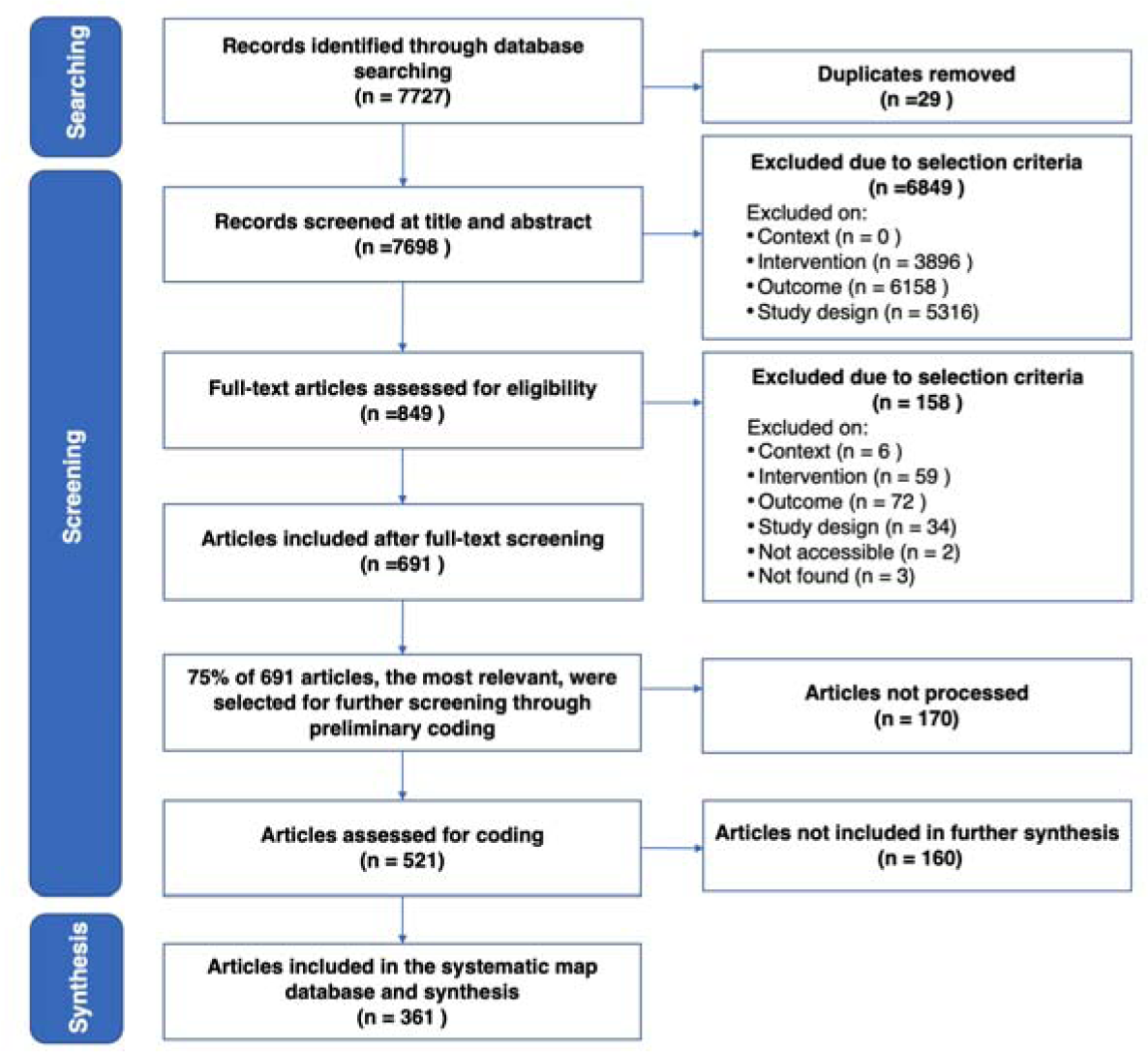
Overview of the selection process of the peer-reviewed publications included in the systematic map database. Flowchart adapted from ROSES flow diagram for systematic maps (Haddaway et al., 2017).

### 2.1 Systematic mapping protocol and the scope of the review

We first established a protocol that describes the systematic mapping methodology, to promote transparency and reduce bias in the review process. The protocol defines the scope of the review based on the CIOS framework (James et al., 2016), which stands for four components of scientific evidence: Context, Intervention, Outcome, and Study type (Table 1). CIOS was used to develop our search strategy and establish the criteria for the inclusion of relevant studies. The Context defines the specific setting where the intervention took place, i.e., urban areas. The Intervention refers to implemented NbS. The Outcome, i.e. the effects of the intervention, included all measured and observed effects related to the NbS. Finally, the Study type was limited to empirical studies, as our analysis targets applied and evaluated NbS. Moreover, the combined assessment of urban challenges, NbS and outcomes required evidence from context-specific cases that reflect complex decision-making and planning. Hence, modelling studies that solely focused on NbS scenarios without empirical data were excluded. The steps of our protocol (search, screening, data coding) are further explained in the following sections. The full protocol is provided in Appendix A.

**Table 1.** Inclusion and exclusion criteria based on the Context, Intervention, Outcome, and Study type (CIOS) framework.

| CIOS element | Inclusion | Exclusion |
| --- | --- | --- |
| Context | Any outdoor environment in urban areas. | Regional, national, and continental context, as well as indoor environment. |
| Intervention | Studies cover the implementation of a nature-based solution, i.e. either specific types of spatial units (e.g., forests, green walls) are changed, or specific actions to alter the spatial units (e.g., revegetation, soil rehabilitation) are implemented. | Interventions that were defined not by any precise feature but only by general terms (e.g., green area, stormwater management systems). Engineered solutions not employing nature. |
| Outcome | Studies report outcomes explicitly related to the effects of nature-based solution interventions (e.g., plant biodiversity, heat reduction). | Studies focused on mechanism/process research and confounding factors (i.e., traffic conditions) without direct evidence on the effects of nature-based solution intervention. |
| Study type | Studies involve empirical evidence (i.e., studies yielding primary data) with a quantitative or qualitative evaluation of the implemented nature-based solution intervention. | Studies on scenario modeling, planning approaches, and conceptual frameworks without providing relevant empirical evidence. |

### 2.2 Search process

Our search strings were based on the Context and Intervention elements of the CIOS framework. For Context, search strings were composed of terms related to urban areas, such as ‘urban’ and ‘city’. For Intervention, a broad set of terms representing approaches related to the NbS concept was applied to capture scientific research that applied NbS-related thinking without necessarily using NbS terminology. To facilitate the development of the term list, we adapted the five categories of ecosystem-based approaches by Cohen-Shacham et al. (2019), to which we added the category ‘nature-based approaches’. We formulated an initial list of specific search terms for each category, thereby also including terms from existing systematic reviews on NbS (Dick et al., 2020; Nesshöver et al., 2017). To reduce bias, terms that focused on specific issues (e.g., climate, disaster, and flood) or ecosystem types (e.g., forest and sustainable drainage) were omitted. Finally, to capture the full breadth of NbS, we reviewed the titles and keywords of 119 review papers on NbS retrieved in August 2021. Terms from those papers that overlapped with or substituted NbS were incorporated across all six categories. Examples include ‘nature-inspired solutions’, ‘sustainable land management’, ‘green spaces’, and ‘blue spaces’. All search strings are provided in Appendix A.

We searched for English language publications listed in the Scopus database, which has a broader and more up-to-date bibliographic coverage than the Web of Science Core Collection. We excluded reviews, proceedings, book chapters, editorials, opinions, and perspectives. We restricted the publication year from 2015 onwards, because the notion of NbS had only been adopted widely in academic research since then (Cohen-Shacham et al., 2016; Maes & Jacobs, 2015). The search was finalized on 25^th^ October 2021 and resulted in 7698 peer-reviewed scientific publications (after removing duplicates).

### 2.3 Screening process

Articles were screened in two stages: first the titles and abstracts, followed by full texts. We used CADIMA, a web tool for conducting and structuring systematic reviews (https://www.cadima.info), to organize the screening process and record selection decisions. A series of eligibility and exclusion decisions based on CIOS was consistently applied to determine the relevance of articles (Table 1). Studies were included in the final dataset only if all CIOS criteria were met. During the screening of titles and abstracts, we refined the criteria for clarity and consistency through two rounds of structured consistency checks. We did so for 80 randomly selected publications in total. For each round, at least three researchers independently screened studies by titles and abstracts and determined the eligibility of papers by scoring all CIOS criteria. Any conflicting scores were resolved through consensus among all authors. This process guided better framing and refinement of the selection criteria. Overall, 89% of the articles were excluded after screening titles and abstracts. Publications with any doubts regarding eligibility were preliminarily retained for the full-text screening.

The full-text screening involved a more precise check on each criterion. After a first iteration of full-text screening, 691 articles were included for further examination. Due to time constraints and the limited relevance of part of these screened articles, we selected the 75% most relevant publications of each publication year to process the full-text screening. Relevance was determined in the Scopus search engine based on the 1) number of hits, 2) significance of the search terms, 3) section the term is mentioned in, 4) position of first occurrence, 5) proximity to other search terms and 6) completeness (Scopus, 2024). In a second iteration, preliminary coding of NbS interventions and their outcomes was done, to more closely examine eligibility based on data collection, measurements, indicators, key results, and interpretation of urban NbS. Through this process an additional 160 publications were excluded, mainly because they lacked direct evidence on the effects of NbS interventions or studies focused on interventions that we did not consider as NbS. Ultimately, after full-text screening, 361 publications were included in the synthesis. Full details of inclusion criteria can be found in Table A4, Appendix A, and the list of exclusion with reasons and included publications in Appendix B.

### 2.4 Data coding strategy

For the included articles, we considered each unique combination of NbS intervention and urban context (location) as a separate case. 547 cases were identified from 361 publications. Two researchers conducted the final data extraction. Overall, 27% of the identified publications were assessed by both researchers, to monitor consistency. The coding was performed manually and documented in FileMaker Pro 19.

For each NbS case, we extracted the following information: 1) geographical locations, 2) NbS, 3) urban challenges, 4) outcomes, and 5) the performance compared to alternative solutions. To identify categories of NbS, urban challenges, and outcomes, we followed an inductive, bottom-up method. Specifically, we proposed the classification based on the detailed data in our database, i.e. without a pre-determined classification.

To establish the classification of NbS types, we first recorded the terms used by the original papers and included a detailed description of the characteristics. These descriptions were then clustered into NbS categories. In this study, **NbS units** refer to spatial units that comprise different elements of natural and built infrastructure in urban areas (e.g., forests, parks and green roofs) (Castellar et al., 2021). **NbS units** were classified based on three features: **vegetation**, **water**, and **engineered**. In addition, a **hybrid feature** was considered as a combination of the three others with no feature dominating. This yielded 16 unit types, in addition to ‘others’ for cases that did not specify the features (Fig. 3). Next, we consider **NbS measures** as actions to manage, restore, or protect existing units, by applying techniques (e.g., replanting and soil rehabilitation) (Castellar et al., 2021). The **NbS measures** were differentiated based on the physical components to which the actions are applied, namely **plants, soil, or abiotic components**. This resulted in eight types of measures, including ‘combined measures’ for cases in which multiple measures were applied (Fig. 3).

**Fig. 3.**
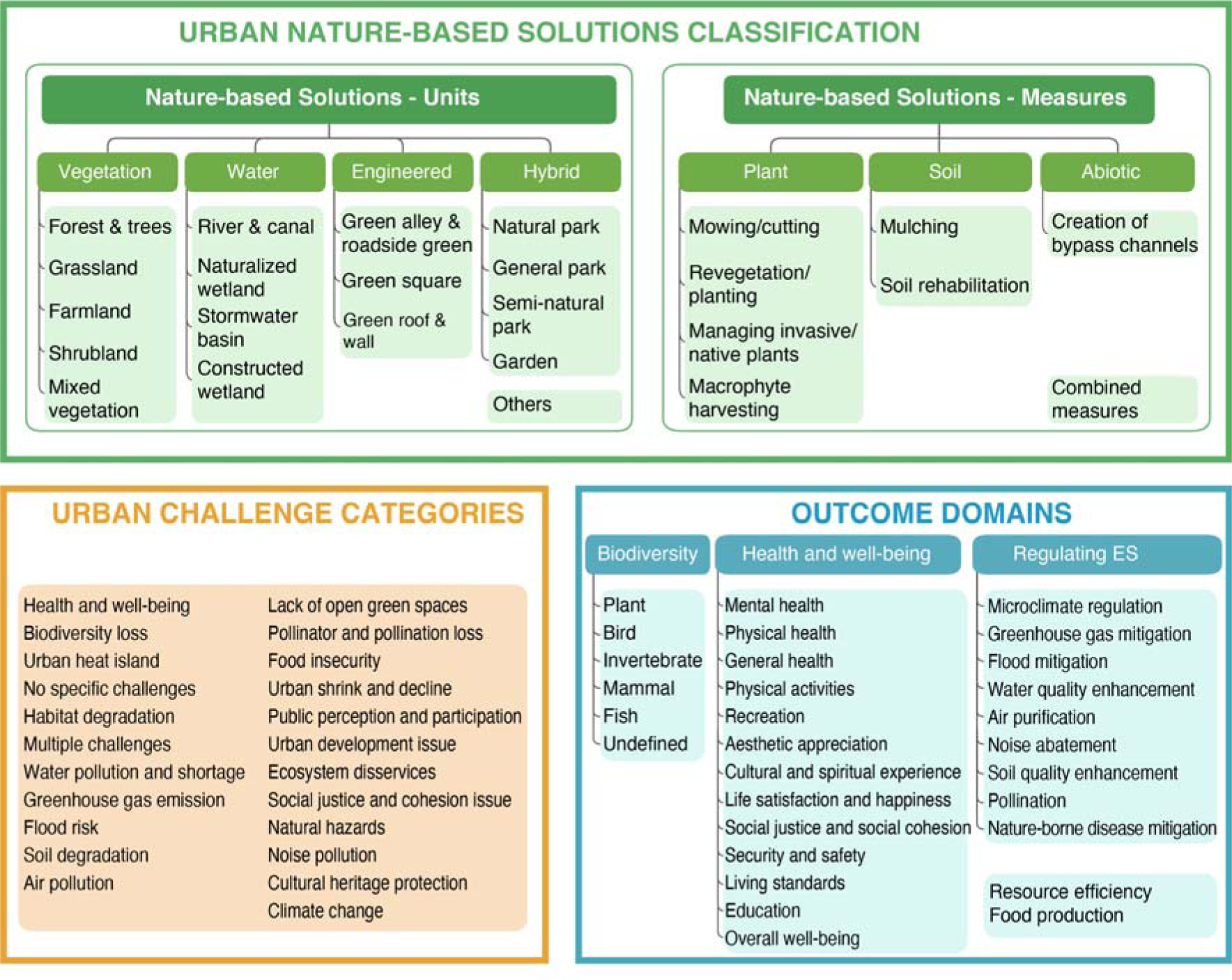
Categories of urban nature-based solutions (NbS), urban challenges and outcomes used in this study.

To map the urban challenges, we adapted a typology of 23 urban challenges from existing categories (Babí Almenar et al., 2021; Dumitru & Wendling, 2021) (Fig. 3). We only recorded an urban challenge if it was stated an issue that could be mitigated by the studied NbS. We also coded cases reporting multiple urban challenges and included a ‘no specific challenges’ category.

To delineate the evaluated outcomes of NbS, we developed a comprehensive list of outcome domains. For each NbS case, we documented all NbS outcomes that had been evaluated using indicators. Domains were based on clusters of similar specific outcomes. For instance, the physical health outcome encompasses outcomes measured by respiratory health conditions, mortality rate, etc. A total of 30 domains were determined (Fig. 3). We categorized these domains into three broad domains: biodiversity, regulating ecosystem services, and health and well-being. None of the outcome domains was found to overlap within or outside the three broad categories.

We considered the performance of NbS as the extent to which an NbS provides an outcome as compared to a non-NbS, i.e., a control with no intervention. We categorized the comparators into control site, before-and-after, green exposure, simulated non-NbS and combination. We recorded the performance as having a positive, negative, neutral or mixed effect. Furthermore, we coded the temporal scale related to these effects, differentiating between a cross-sectional effect (i.e., evaluated at a single time point) and an effect over time (i.e., evaluated across various time points). We also documented whether the NbS case reported multiple outcomes for further synthesis of multifunctional NbS. Full details of all the coding categories and their definitions can be found in Appendix A.

### 2.5 Data analysis and visualization

We mapped the geographical locations of identified urban NbS and then analyzed the distribution of the evidence regarding NbS types, urban challenge categories, and outcome domains (547 cases with 799 outcomes in total) (Section 3.1). Subsequently, the coded components regarding urban challenges, NbS, outcomes and performances were used to systematically analyze the effectiveness of NbS to address urban challenges. Sankey diagrams were created to identify and visualize the linkages between urban challenges, NbS, and outcomes (Section 3.2). This analysis included only cases of the most frequently reported urban challenge categories (reported in over 15 cases) and outcome domains (idem). A total of 462 cases were considered, involving 630 outcomes. The performance of NbS was synthesized considering outcome domains, NbS unit types, comparison types and temporal scales (Section 3.3). We also analyzed the cases that reported potential synergies (multiple positive effects) and trade-offs (positive and negative effects together). This analysis involved 370 outcomes and their non-NbS comparators, based on 280 cases. Finally, we conducted a co-occurrence analysis between outcomes, for the 133 cases reporting multiple outcomes (Section 3.4).

All data analyses were conducted in R (v.4.2.2) with RStudio (v.2022.07.2+576), except the global distribution map (Fig. 4A), which was created using ArcGIS 10.6. For the data visualization, the following R libraries were used: ggplot2 (Figs. 4B-F, 7, and 8) and networkD3 (Figs. 5 and 6). Additional analyses that supplement the main results can be found in Appendix C.

**Fig. 4.**
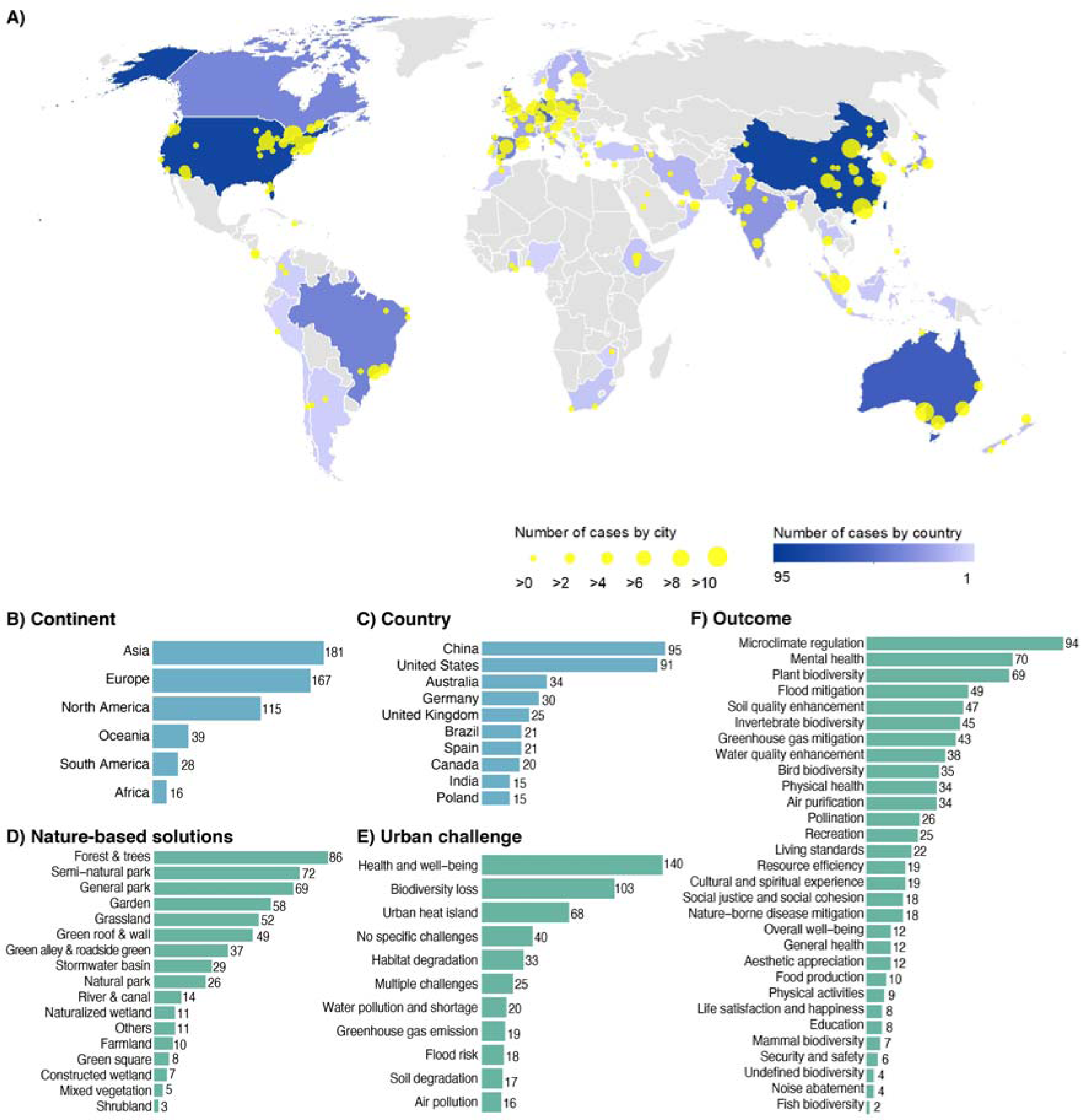
Overview of the systematic mapping output, with (A) geographical distribution of cases, (B) the number of cases per continent, (C) the 10 countries with the highest number of cases, (D) the number of cases per nature-based solution, (E) the number of cases of the 11 most-reported urban challenges, and (F) the number of assessments per outcome domain.

**Fig. 5.**
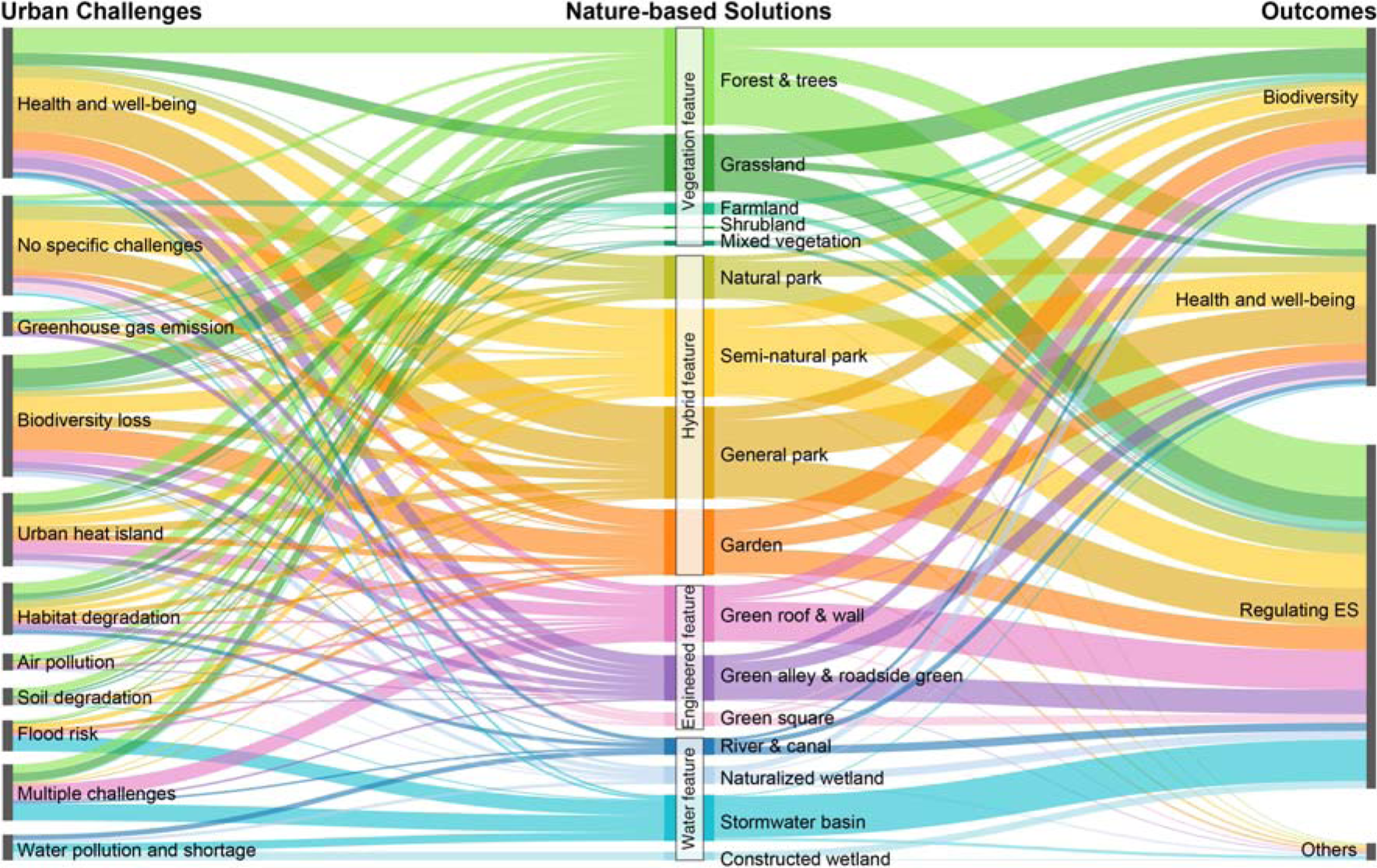
Sankey diagram relating 16 nature-based solutions (NbS) unit types and 4 features (middle of the diagram) to the 11 most reported urban challenges (left) and 3 broad outcome domains (right). The data used in the analysis contains 630 outcomes related to 462 NbS cases. The thickness of each band corresponds to the number of cases. NbS types were arranged and color-coded according to features: those in green correspond to the vegetation feature, those in yellow to the hybrid feature, those in purple to the engineered feature, and those in blue to the water feature.

**Fig. 6.**
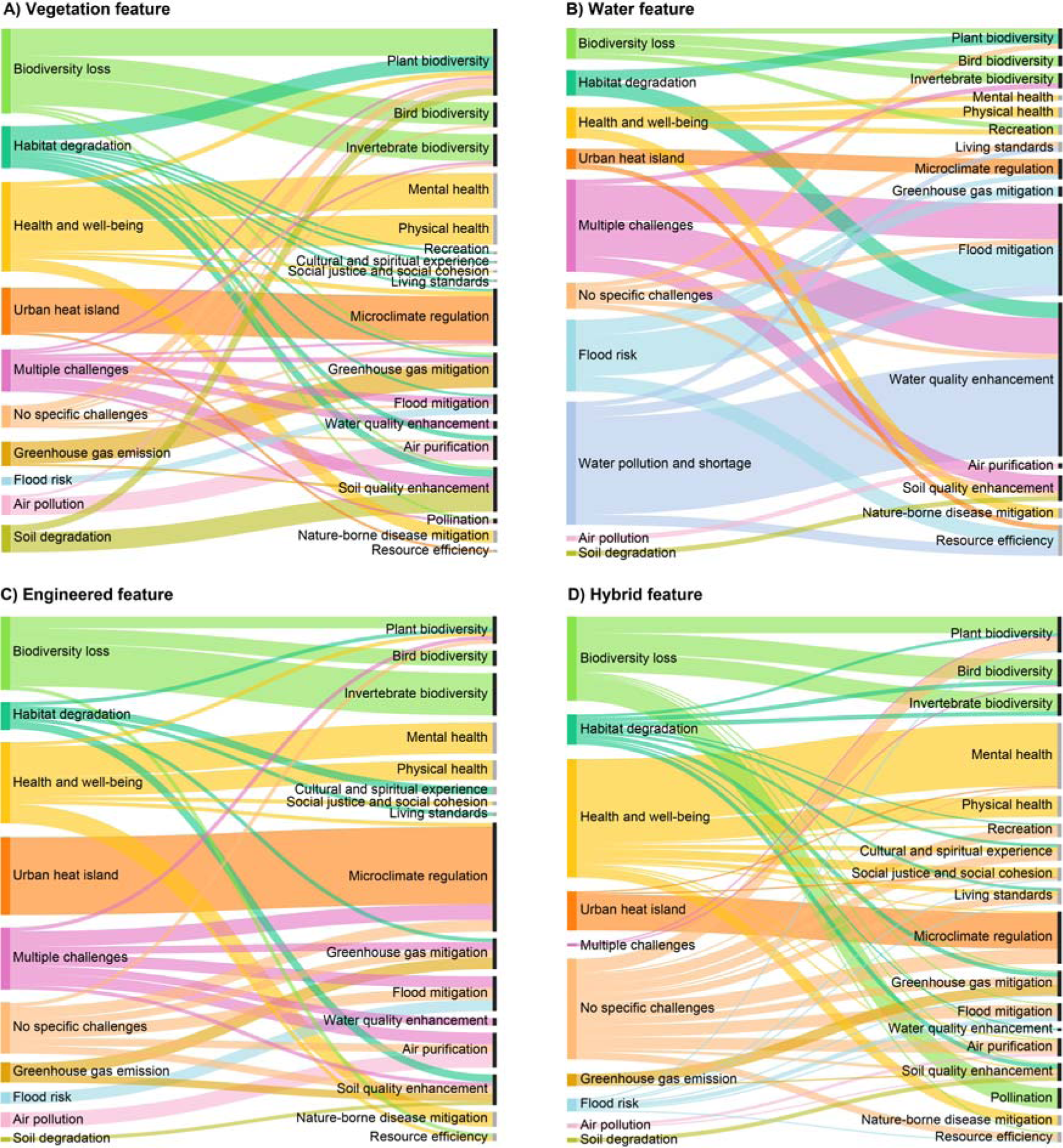
The linkages between urban challenges (left) and outcomes (right) for 4 features of NbS: A) Vegetation feature, including 132 urban challenges and 164 outcomes; B) Water feature, including 58 urban challenges and 84 outcomes; C) Engineered feature, including 79 urban challenges and 109 outcomes; and D) Hybrid feature, including 163 urban challenges and 273 outcomes. Outcomes (right) are arranged by biodiversity (black nodes), health and well-being (grey nodes), regulating ecosystem services (black nodes), and others (grey nodes).

## 3. Results

### 3.1 Geographical distribution, nature-based solutions, urban challenges, and outcomes

#### 3.1.1 Geographical distribution

The majority of cases were from Asia, Europe, and North America, together covering 84% of all the cases (Figs. 4A and 4B). Evidence was reported from 56 countries, with China (17%) and USA (16%) providing substantially more cases than other countries (Figs. 4A and 4C). More than half of the Asian cases are from China (53%) and 80% of North American cases are from USA.

Evidence was reported from 196 unique cities, and 21 additional cases did not specify the city. A large number of studies was done in Beijing (27), New York (22), and Singapore (13). 99 cities only had a single case.

#### 3.1.2 Nature-based solutions

Among 547 NbS cases identified, NbS with hybrid features, such as parks and gardens, were most frequently reported (41%). NbS with vegetation features were the second most reported (29%), while NbS with engineered and water features were less represented (17% and 11%, respectively). Forest & trees were the most studied NbS unit type (16%), followed by semi-natural park and general park (13% each), garden (11%), grassland (10%), and green roof & wall (9%) (Fig. 4D). Only in 11% of the cases were measures applied within units (n=62). These measures mostly involved plants (56%) or soil (42%) and to a limited extent abiotic components (2%). A combination of measures was observed 29% of 62 cases. Mowing/cutting was the second most reported measure type (28%), which was mainly applied in grasslands (13 out of 16 cases). Combined measures were most related to river and canal NbS (8 out of 18 cases).

#### 3.1.3 Urban challenges

Overall, 88% of the cases reported an individual urban challenge and 5% of the studies reported 2 or 3 different urban challenges. In 7% of the cases, no clear urban challenge was reported. The most often reported urban challenges include health and well-being (26%), biodiversity loss (19%), and urban heat island (12%). 91% of the cases were found to fall within the 11 major challenge categories, each represented by more than 15 cases (Fig. 4E).

#### 3.1.4 Outcomes

Most of the outcomes (Fig. 4F) were related to regulating ES (44%), followed by health and well-being (32%) and biodiversity (20%). Of the 353 regulating ES outcomes, microclimate regulation (i.e., heat reduction) was most reported (27%), followed by flood mitigation (14%), soil quality enhancement (13%), greenhouse gas mitigation (12%), and water quality enhancement (11%). Of the 255 outcomes of health and well-being, most were related to mental health (27%) and physical health (13%). Few outcome assessments involved social well-being outcomes, such as aesthetic appreciation, life satisfaction and happiness, and security and safety (each around 1%). Out of the 162 biodiversity outcomes, most focused on plants (43%), invertebrates (28%), and birds (22%), while few assessed mammals (4%), undefined biodiversity (2%), and fish (1%).

### 3.2 Links between urban challenges, nature-based solutions, and outcomes

#### 3.2.1 Nature-based solutions connecting urban challenges and outcomes

Most NbS were linked to all three overarching broad domains of outcomes: health and well-being, regulating ES, and biodiversity (Fig. 5). The exceptions were stormwater basins and constructed wetlands, which did not report any biodiversity outcomes, and constructed wetlands, farmlands, shrublands, and mixed vegetation, which did not report any health and well-being outcomes.

In general, forests & trees, semi-natural parks, general parks, and grasslands are the NbS types with the highest number of associated urban challenges (Fig. 5). Among NbS with vegetation features, most links involve forests & trees addressing health and well-being (24), and grasslands addressing biodiversity loss (18). Of NbS with hybrid features, many cases link general parks to health and well-being (33), and gardens to biodiversity loss (23). NbS with engineered features most commonly involve links between green roofs & walls and the urban heat island (12), and green alleys & roadside green addressing health and well-being. Among NbS with water features, stormwater basins were commonly linked to flood risk (14), while constructed wetlands and stormwater basins, tackled water pollution and shortage (8 each). A combination of urban challenges was most often addressed by stormwater basins (16) and green roofs & walls (14). The highest number of links to no specific urban challenges was related to semi-natural parks (24), general parks (23), and natural parks (14).

#### 3.2.2 Links between urban challenges and outcomes

Linkages between outcomes and urban challenges differ per NbS unit feature, especially between water and the three other features (Fig. 6). The most frequently mentioned urban challenges, i.e. health and well-being, biodiversity loss, and the urban heat island, exhibited the highest numbers of links to outcomes in all features, apart from water. Within the 84 links associated with the water feature, water pollution and shortage (36%), and flood risk (12%) showed the highest number of links to outcomes. Moreover, water is the only feature with links to the challenge of water pollution and shortage, and without links to the challenge of greenhouse gas emission.

In most cases, an urban challenge was linked to the associated outcome that would be most intuitive (Fig 6). For instance, the urban heat island challenge was most often linked to the microclimate regulation outcome (65 of the 69 total links to the urban challenge), greenhouse gas emission to greenhouse gas mitigation (19 of 23 links), and biodiversity loss to the biodiversity of invertebrates (36 of 115), birds (29) and plants (26). The multifunctionality of NbS was shown by the fact that additional outcomes were linked to a specific urban challenge. These include the links between health and well-being challenge and regulating ES outcomes (e.g., microclimate regulation), but also biodiversity outcomes (e.g., plant biodiversity). In addition, the biodiversity loss challenge was often linked to additional regulating ES outcomes (e.g., pollination).

Our analysis also shows disconnections between specific outcomes and urban challenges. First, 7% of cases do not report an urban challenge. As a result, 16% of the outcomes have no link to a specific urban challenge (Fig. 6). The majority of disconnections occurring within the hybrid feature (67 of 94) link to the ‘no specific challenge’ category (Fig. 6). Furthermore, many social well-being outcomes lack a connection to an urban challenge. Among those, cases involving recreation (7 out of 12 links missing), cultural and spiritual experience (6 out of 14), and living standards (6 out of 15) most often do not state specific urban challenges. In addition, social well-being outcomes were not connected to the urban challenge of health and well-being. For example, cultural and spiritual experience, living standards, and recreation outcomes provided by vegetation features were only related to the challenge of habitat degradation (Fig. 6A).

### 3.3 Performance compared to non-nature-based solutions

Of the 370 performance assessments that were compared to a non-NbS, 79% reported a positive effect, 5% a negative effect, 8% a mixed effect, and 8% a neutral effect (Fig 7). For microclimate mitigation, the performance was positive in 91% of the 65 cases (Fig. 7A). Invertebrate biodiversity exhibited the highest proportion of negative effects (50%), of which most are attributed to mowing or cutting measures in grasslands. Additionally, negative effects were reported for 13% of physical health outcomes, all of which were assessed through comparison of green exposure. Performance comparisons were available for all outcome domains except aesthetic appreciation and education.

**Fig. 7.**
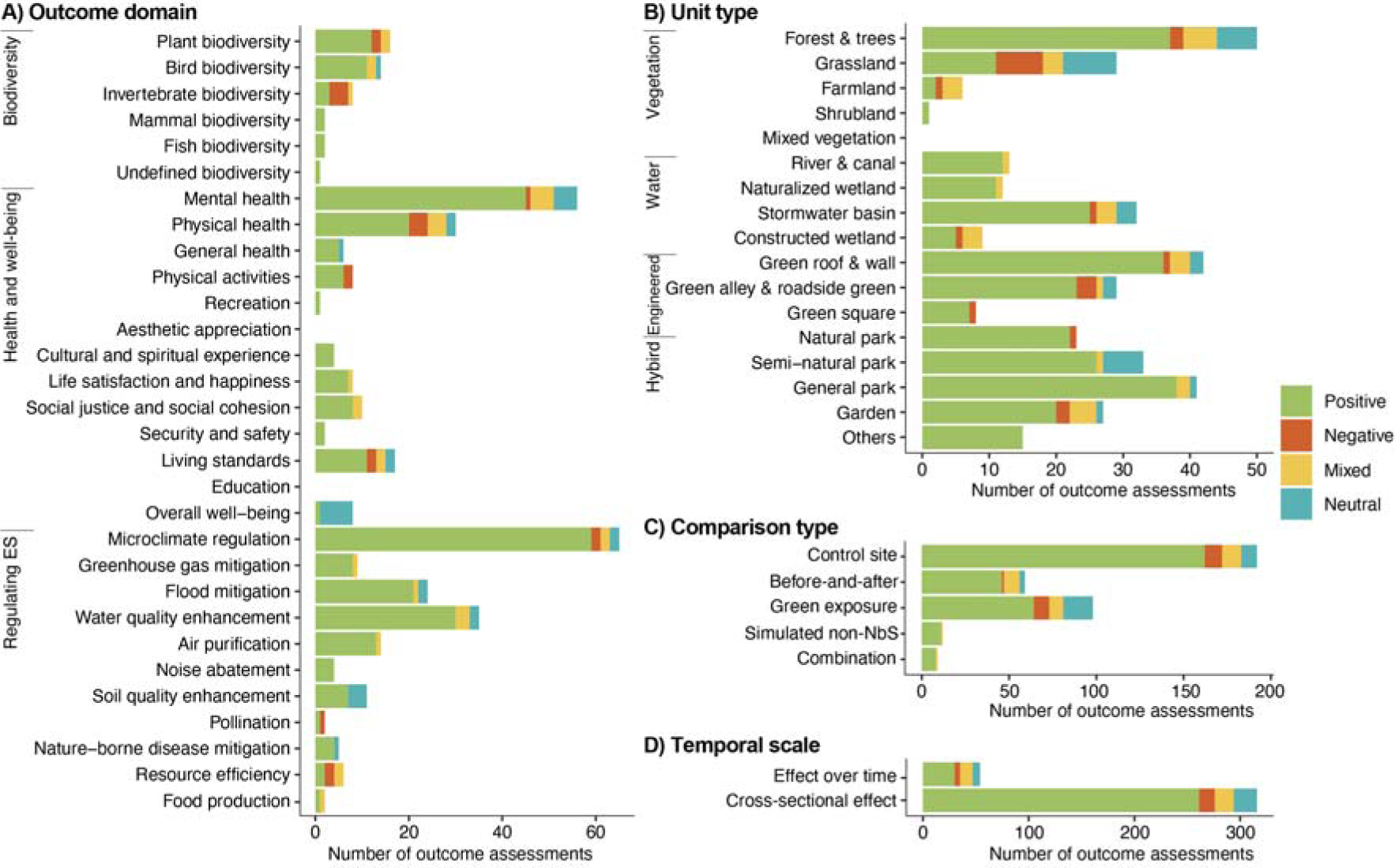
Performance assessed by comparing NbS outcomes to those of a non-NbS. The performance (positive, negative, mixed and neutral effects) is categorized by outcome domain (A), unit type (B), comparison type (C), and temporal scale (D). The analysis was based on 370 outcomes.

Among NbS types, natural parks, general parks, rivers & canals, naturalized wetlands and green roofs & walls exhibited a higher proportion of positive effects than average (Fig. 7B). Negative effects were less prevalent in these unit types (all below the 5% for all outcomes combined). Negative effects were most frequently reported for grasslands (24%), farmlands (17%), green squares (12%), and green alleys & roadside green (10%). Notably, 10 out of 12 positive effects reported from rivers & canals were related to measures taken within this NbS type, mostly involving combined measures (8). Out of 22 negative effects, 5 were associated with measures, primarily linked to mowing/cutting (3).

Regarding the temporal scale of performance, 85% of the 370 reported effects were cross-sectional, while 15% involved effects over time (Fig. 7D). Cross-sectional effects were mostly positive (83%), while effects over time demonstrated comparatively fewer positive effects (56%). A high percentage of plant biodiversity outcomes were reported as effects over time (65%).

The performance of NbS with multiple outcomes could be evaluated against non-NbS in 47% of the 133 cases. Of these cases, 32% provided multiple positive effects, indicating the potential for synergies between NbS outcomes. These cases primarily involved general parks (4) and stormwater basins (3). General parks were found to provide multiple benefits such as physical health, mental health, and life satisfaction and happiness. Stormwater basins addressed challenges of water pollution and shortage, or flood risk, by providing multiple positive effects such as flood mitigation, water quality enhancement, and greenhouse gas mitigation. Only one case indicated a potential trade-off, where a constructed wetland had a positive effect on water quality enhancement (nitrogen removal) and a negative effect on resource efficiency (increasing water consumption). A full list of cases reporting potential synergies is available in Appendix C.

### 3.4 Evidence of multiple outcomes and co-occurrence analysis of outcomes

Multiple outcomes were documented in 24% of the NbS cases (133), involving 387 outcomes in total. Among these cases, 59% reported two outcomes and 41% reported three or more outcomes. One-third reported outcomes related to more than one broad domain, i.e. health and well-being, regulating ES, or biodiversity. Most cases of multiple outcomes were reported for forests & trees (20), followed by semi-natural parks, general parks (each 13), and gardens (12).

The highest frequencies of co-occurrences were found between two outcomes within the same broad domain (Fig. 8). For instance, flood mitigation was most frequently associated with three other regulating ES: water quality enhancement, microclimate mitigation, and air purification (14 each). Likewise, mental health co-occurred most often with physical health (14). Our analysis also revealed that outcomes within the same broad domains commonly coincide. Notably, flood mitigation co-occurs the most with other outcomes (100), followed by microclimate regulation (88) and air purification (76). Regulating ES outcomes frequently co-occurred with health and well-being outcomes, such as microclimate regulation with recreation, greenhouse gas mitigation with cultural and spiritual experience, and air purification with aesthetic appreciation (7 each). Of the biodiversity outcomes, only plant biodiversity co-occurred commonly with regulating ES, such as greenhouse gas mitigation and flood mitigation (7 each). Few other cases of co-occurrence were observed between health and well-being outcomes and biodiversity outcomes.

**Fig. 8.**
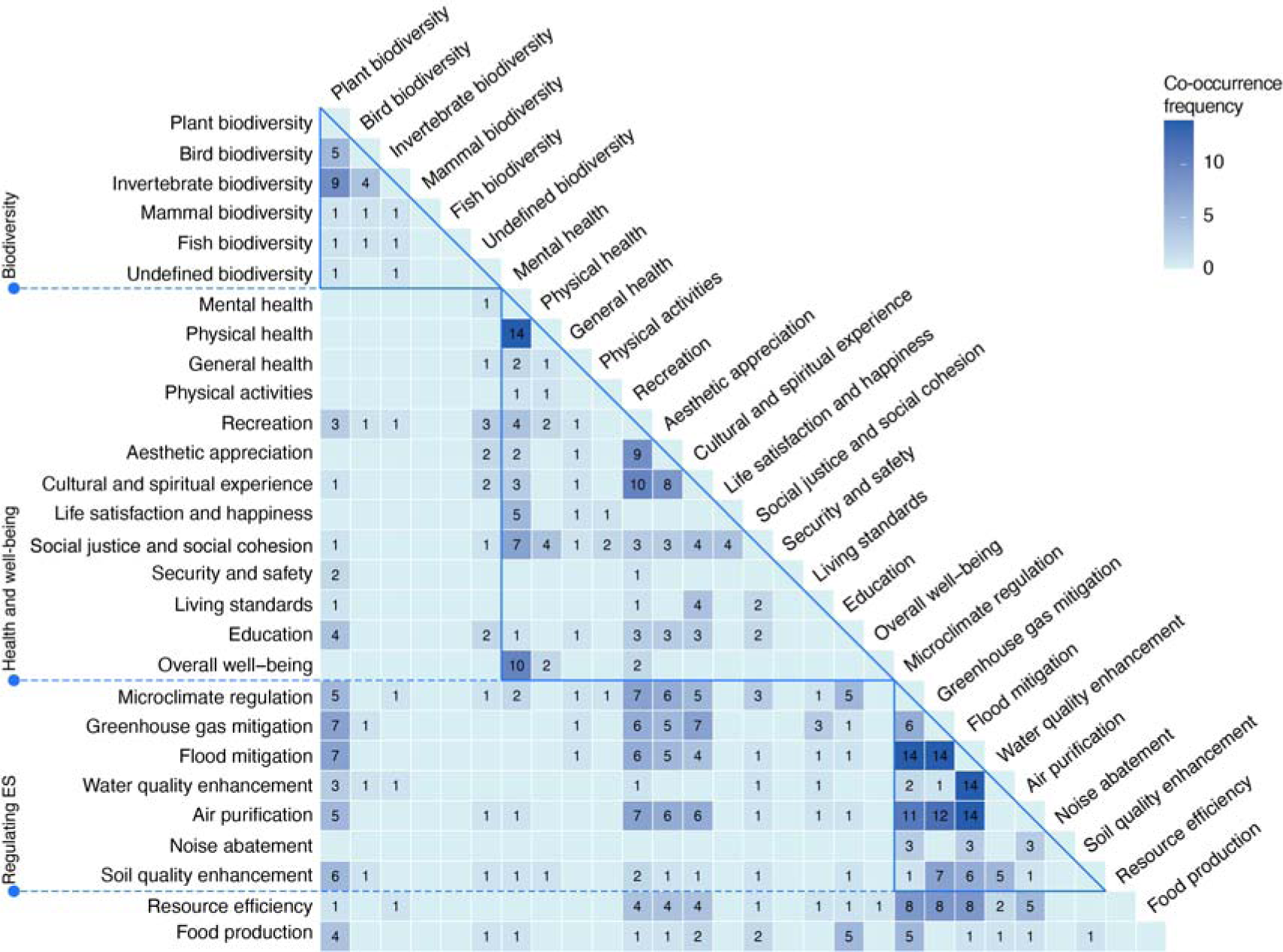
Co-occurrence analysis between outcome domains of nature-based solutions. The data includes 387 outcomes generated from 133 cases containing multiple outcomes. The heatmap visualizes the 28 outcome domains that co-occur with others. Outcome domains are arranged by the broad domains of biodiversity, health and social well-being, regulating ES, and others.

## 4. Discussion

Through the combined assessment of NbS types, urban challenges, and outcomes, this study provides a foundation for a more comprehensive understanding of NbS effectiveness in urban areas. Our assessment provides three critical assessments of effectiveness, including how urban challenges are addressed by NbS with associated outcomes, their performance compared to alternative solutions and the potential to provide multiple outcomes. To the best of our knowledge, this is the first global systematic mapping that combines these different components of NbS effectiveness, especially for urban systems. Our research addresses the call for a better evidence base of NbS effectiveness to address urban challenges (Fang et al., 2024; Johnson et al., 2022).

Our database complements the existing knowledge base by synthesizing evidence from 547 urban NbS cases in 197 cities, linking NbS outcomes to urban challenges and thus providing evidence of effectiveness. In this way, our systematic analysis helps policymakers and urban planners with information tailored to their specific interests. For instance, our analysis presents a list of urban NbS types and a range of associated outcomes that they can prioritize to address urban challenges or the growing demand for multifunctional NbS. Moreover, such evidence-based research can help mainstream NbS in urban policy by providing guidance on the most suitable set of NbS types that can effectively contribute to addressing specific urban challenges (Cohen-Shacham et al., 2019; Frantzeskaki et al., 2019).

Following recent efforts to align NbS with ES in urban areas (Remme et al., 2024), we provide evidence on outcomes of urban NbS linked to ES, as well as broader benefits. We provide a fine-scale assessment of specific urban green interventions, which is rare in ES research (Veerkamp et al., 2021). Compared to previous reviews that assessed the linkages between urban sustainability challenges, NbS, and ES (Babí Almenar et al., 2021; Fang et al., 2024), our study employed a method that allowed for a more comprehensive assessment beyond ES. This particularly involves a broader coverage of outcomes related to human health and well-being, and biodiversity conservation. We also advanced beyond the examination of the linkages between these outcomes and urban challenges, by further assessing the outcome performance and cooccurrence of multiple outcomes.

To assess NbS effectiveness in a balanced manner, we considered a broad range of challenges and outcomes related to biodiversity, well-being, and regulating ES. We did not start from a predetermined list of outcomes or challenges. This resulted in a more holistic assessment of linkages between challenges, NbS, and outcomes, compared to the existing body of synthesis work (Kabisch et al., 2017; Villamayor-Tomas et al., 2024). Our analysis provides more insight into a broader range of urban challenges, a wide, yet well-defined, set of NbS, and a large variety of related outcomes. Our broader also scope allowed us to explore the multifunctional potential of urban NbS with 132 empirical cases involving multiple outcomes. This facilitated a novel synthesis of intersections between evaluated outcomes for both biodiversity and human well-being, which helps advance the knowledge on a broad definition of effectiveness beyond assessing if specific targets are met.

Our search terms and inclusion criteria allowed us to look beyond research focusing only on the term ‘nature-based solutions’. This offered an overarching perspective on a swiftly developing concept, without disregarding empirical studies in parts of the world or research fields where the concept is called or conceptualized differently. The assessment was conducted on the most relevant 75% of the identified articles after screening (Fig. 2). We conducted a quick examination of the excluded 25% of articles, coding continental distribution, urban challenge categories, NbS unit types, and outcome domains. This examination confirmed that processing the complete list of articles would not have had a discernible impact on our findings (Appendix A).

In the subsequent sections, we first reflect on the urban NbS classification employed in our research. Next, we discuss the key findings of the three aspects related to the effectiveness of urban NbS, and finally their implications for research and practice, including important evidence gaps and related opportunities for future synthesis and research.

### 4.1 Classifying nature-based solutions in the urban context

NbS are defined, classified, and interpreted differently throughout the literature (Dorst et al., 2019). Castellar et al. (2021) and Babí Almenar et al. (2021) proposed classifications of NbS in urban contexts, further modifying the three types of NbS defined by Eggermont et al. (2015). Castellar et al. (2021) provided a common conceptualization of NbS by consolidating the knowledge from European projects to facilitate NbS uptake. Babí Almenar et al. (2021) based their classification on ES assessments to facilitate connection between ES and NbS. We built on these classifications with the aim to facilitate the evaluation of effectiveness of urban NbS in a global context. We thus developed a classification rooted in the scientific literature of NbS outcome evaluation and its linkage with urban challenges.

In our classification, NbS types involve a detailed examination of physical features, both natural and built components. During the conceptualization and execution of our study, many iterations and discussions were essential to help interpret and classify different aspects of the NbS concept. This classification is intrinsically linked to the primary point that involves the interpretation of what constitutes ‘nature-based’. Even though there is no consensus on what constitutes nature (Kotsila et al., 2021), our operationalization was in line with the CBD’s definition of an ecosystem, i.e. consisting of various natural elements such as plants, animals, soil, and water, and their interactions (United Nations, 1992). Similar to Castellar et al. (2021), we considered that vegetation plays an important role in characterizing an NbS. The growth of plants embodies a living ecosystem capable of supporting biodiversity, a fundamental tenet to NbS (Key et al., 2022). Following this logic, even bio-engineered solutions that mimic nature, such as permeable pavements, can be defined as NbS, as long as they provide physical space for vegetation to grow.

Another point of attention relates to when nature can be considered an intervention and, importantly, a ‘solution’. We argue that intentionality is crucial to qualify an NbS, i.e. units or measures involving natural elements being implemented as an intervention to tackle a challenge or achieve a goal. This includes an intentional decision to leave an area untouched in order to achieve particular goals. Cities contain informal green spaces and abandoned areas that are neglected or not purposely managed and maintained, e.g. emerging forest patches in abandoned areas and engineered infrastructure with spontaneous plant growth (Huang et al., 2019; Kowarik et al., 2019). However, these ‘interventions’ can only constitute an NbS if such spaces are managed by ‘doing nothing’ rather than building in them (Sikorski et al., 2021), and such decisions are coupled with the identified challenges such as preserving biodiversity.

Our classification couples measures to physical NbS units and therefore facilitates the consideration of management approaches in NbS urban planning. Up until now, interventions involving protection, management, and restoration approaches were considered as categories of NbS in existing classifications, along with created ecosystems (Castellar et al., 2021; Eggermont et al., 2015). Yet, these categories have proven challenging to distinguish as separate solutions, or often intangible and hard to apply in urban contexts, as shown by the near-absence of management approaches in the review of urban ES literature (Babí Almenar et al., 2021). In our database, most cases involved NbS units without specific management or measures. Only 11% of the cases involved a measure applied to an NbS unit. However, the cases that did couple measures to physical units provide valuable insights for urban planners on how to effectively manage the existing ecosystems. For example, research indicates that reducing mowing frequency in grasslands can contribute to bee conservation. Such information holds significance for cities that have limited space for creating new ecosystems.

### 4.2 Linking outcomes of nature-based solutions to urban challenges

Understanding urban challenges targeted by NbS and the outcomes generated by them is crucial for assessing the effectiveness of NbS. Our results confirm that the majority of cases (507 of 547) explicitly link NbS to specific urban challenges. Cases for which this link was missing mostly related to studies focused on ES assessment and economic valuation (e.g., Chen, Wang, Ni, Zhang, & Xia, 2020; Hoover, Price, & Hopton, 2020). In our dataset, the most frequently reported urban challenges include health and well-being, biodiversity loss, and the urban heat island. We established that tree-based NbS (forests, street trees) and parks most often target health and well-being challenges, whereas grasslands and gardens involve challenges of biodiversity loss. Compared to urban NbS reviews by Fang et al. (2024), our study includes a higher coverage of challenges related to biodiversity loss and human health issues. Babí Almenar et al. (2021) included more challenges related to circular economy and material/resources management, emphasized sustainability and resilience more strongly, and included grey literature. Fang et al. (2024) focused on research explicitly using the term NbS and found a primary focus on climate change. Our broader search scope, using a wider array of NbS-related search terms, captured a more diverse set of challenges.

Although reported outcomes could often be linked to challenges, their frequencies follow different patterns. The most frequently reported outcomes involve regulating ES, followed by health and well-being and biodiversity. Especially microclimate regulation, mental health, and plant and invertebrate biodiversity were commonly reported outcomes. Although they partly match the key outcomes commonly associated with NbS (Dick et al., 2020; Key et al., 2022), especially well-being outcomes such as safety and security, life satisfaction and happiness, and aesthetic appreciation were underrepresented in our data. Such data is often reported on larger scales (i.e. nation-or citywide) and not in relation to a particular spatial unit. Furthermore, our findings suggest that multiple outcomes were provided by urban NbS to address specific urban challenges. We also note that the NbS cases were limited in terms of interdisciplinary approaches, with few studies measuring outcomes related to biodiversity and human well-being in combination.

Our findings suggest that, aside from human health, other well-being dimensions have not been adequately addressed in empirical studies on established NbS. Within our database, the health and well-being challenge category was primarily focused on human health issues, with few cases addressing other well-being dimensions such as education, social justice and social cohesion, and living standards. Similarly, physical and mental health outcomes dominated the health and well-being domain, in line with Dick et al. (2020). Furthermore, our results reveal that well-being outcomes other than human health are rarely linked to health and well-being challenges, but rather to other challenges, such as flood risk. These well-being outcomes are often included in NbS research as secondary outcomes of NbS, while primarily addressing other challenges. Although well-being dimensions like social justice have been stressed in studies on urban NbS policy, planning, and governance (Haase et al., 2017; Sekulova et al., 2021), solid evidence of the contribution of NbS to improve the living conditions of urban dwellers and foster social inclusiveness is still needed (Haase et al., 2017). To obtain more comprehensive evidence of NbS effectiveness in line with other entries in our database, we advocate for further empirical studies related to implemented NbS that encompass a broader range of well-being dimensions beyond human health. In addition, the inclusion of empirically grounded studies on preferences and responses to NbS or alternative scenarios (e.g., through surveys) could provide broader insights.

### 4.3 Assessing the performance of nature-based-solutions and ways forward

The performance assessment of NbS compared to non-NbS showed that 79% of the outcomes were positive. This suggests that NbS can be effective solutions compared to constructed or engineered solutions, in line with previous literature (Acreman et al., 2021; Chausson et al., 2020; Kabisch et al., 2017). The finding could also be indicative of a tendency to focus primarily on positive effects in academic publications (Mlinarić et al., 2017). Furthermore, consistent comparisons for some NbS are challenging, due to the large variation in outcomes considered. Microclimate regulation and mental health outcomes were often involved in comparative studies (69% of 94 outcomes and 80% of 70 outcomes respectively). Conversely, biodiversity-related, pollination, greenhouse gas mitigation, and soil quality enhancement outcomes were not often included in comparative studies (25% or less of reported outcomes). For the latter topics, the likely explanation is that the non-NbS comparators do not have the necessary biophysical attributes to make useful comparisons.

Also, education and aesthetic appreciation have no comparative studies. Those topics generally rely on large-scale social surveys rather than city-specific empirical studies. Conducting comparisons of NbS outcomes, while controlling for other variables within the survey, would require additional efforts. To make the outcomes more transferable to urban planners, more comparisons between NbS and non-NbS control are needed, especially for outcomes based on biophysical measurements or related to social well-being.

Our analysis primarily examines the general directionality of outcomes summarized by binary situations (i.e., NbS versus non-NbS controls) to assess general effectiveness, with a strong focus on (bio)physical performance. However, further dimensions of effectiveness, such as the extent and longevity of the performance, remain unclear. For such analyses, a more detailed quantitative synthesis is needed that analyzes the methodology of monitoring and evaluation of each case. Our database revealed a large heterogeneity in quantified outcomes of urban NbS and the variety of methods applied to evaluate their performance, such as comparators and indicators applied. This reduced comparability across studies and resulted in our approach to effectiveness assessment. Moreover, broader aspects of effectiveness, such as economic viability, compliance with legal frameworks, or influence on long-term human behavior could also be incorporated into effectiveness assessments (Seddon et al., 2020; Sowińska-Świerkosz & García, 2021). These aspects were, however, beyond the scope of our current study. Here, we suggest directions for future studies to provide insights into deeper dimensions of effectiveness.

For future synthesis research, a focus on quantifying spatial and temporal effects of NbS on outcomes will help improve the granularity of relationships between challenges, NbS, and outcomes (Remme et al., 2024). First, to quantify NbS performance across different spatial scales, further examination of the physical dynamics underlying the delivery pathways of these benefits, such as green exposure, heat fluxes, and water flows, is required (Kumar et al., 2021; Raymond, Berry, et al., 2017). For example, the benefits of NbS are scale-dependent, whether it be by the extent of impact (e.g., airshed, watershed) or proximity to receptors (e.g., noise, health) (Hutchins et al., 2021).

Second, further temporal assessment needs to focus on the required time of NbS to become effective (Sowińska-Świerkosz & García, 2021). Our synthesis forms the initial step in evaluating the temporal scale of the performance. We distinguish between ‘snapshot’ evaluations at a specific time and those assessing multiple time points. However, the specific length of time for a particular intervention to become effective is not widely documented in the studies in our database or other analyses (Dick et al., 2020; Kumar et al., 2021). We call for clear reporting on several temporal aspects in all NbS studies. First, the starting time and, if relevant, duration of NbS should be recorded. Additionally, assessing the longevity of the performance requires post-intervention monitoring. This can be facilitated by a rigorous framework that incorporates the monitoring process, including techniques, available data for specific indicators, and time-specific data collection frequencies (Kumar et al., 2021).

Further research is recommended to more comprehensively synthesize potential differences in performance across different NbS units and measures. While we recorded comparisons to other NbS types, further synthesis was hindered by the use of multiple heterogeneous NbS comparators without specifying outcomes for each, as also noted by Dick et al. (2020). A recent review provided a possible approach to generalize the comparative results by focusing on specific pairs of NbS types, such as forest and park (Beute et al., 2023). Their results were mixed due to heterogeneity in NbS characteristics and research design. Quantifying outcomes among different NbS types requires meticulous recording of NbS characteristics, including variations in management regimes and morphology variables such as size, tree coverage and plant composition.

### 4.4 Multifunctionality of nature-based solutions

To fully assess the effectiveness of NbS, the synergies and trade-offs between their multiple outcomes need to be understood. The evidence base needs to be further developed, although some relevant insights exist. For example, an integrated evaluation has shown that forest plantations as NbS can provide job opportunities while revitalizing degraded forests and mitigating fragmentation (Lemgruber et al., 2021). Regarding trade-offs, a rooftop farm was implemented as an NbS for ensuring food security in highly dense urban areas, but the life cycle cost increased compared to using a flat roof (Kim et al., 2018). Such evidence is important to understand the interactions between outcomes related to social, ecological, and technical dimensions of complex urban systems underlying the effectiveness of NbS (McPhearson et al., 2022). For example, extensively managed lawns can foster plant species richness but represent high fire risk (Winkler et al., 2021), which shows that management choices in the technical dimension can influence the impacts on ecological and social dimensions. Understanding the interactions can improve urban planning and management choices to maximize synergies and avoid significant trade-offs, to ensure more effective and sustainable NbS (Gómez Martín et al., 2020; Raymond, Frantzeskaki, et al., 2017).

The scarcity of studies evaluating the co-occurrence of outcomes across different broad domains highlights the limited involvement of interdisciplinary approaches for NbS evaluation. Data availability to calculate an integral set of indicators is crucial for the integrative evaluation of an NbS project (Sowińska-Świerkosz & García, 2021), but varying technical expertise needed to measure different aspects of outcomes can be a constraint (Key et al., 2022). Therefore, it is unsurprising that we find the highest number of co-occurrences between reported outcomes within single domains, such as mental health and physical health, since they involve overlapping disciplinary expertise. Recent efforts to develop robust NbS evaluation frameworks, integrating selected indicators across multiple outcomes, present a big step forward (Raymond, Frantzeskaki, et al., 2017; Watkin et al., 2019). However, further development is essential through more empirical studies that apply these frameworks and facilitate refinement by incorporating site-specific feedback from the field (Colléony & Shwartz, 2019; Watkin et al., 2019). Moreover, more theoretical work is required to elucidate the pathways connecting various types of NbS to multiple outcomes, particularly focusing on the intermediate mechanisms linking NbS characteristics to multiple outcomes (Dumitru et al., 2020).

## 5. Conclusions

NbS have been implemented without systematic scientific evidence of their effectiveness concerning multiple outcomes and targeted urban challenges. Based on 547 empirical case studies identified from scientific literature studying NbS outcomes in cities worldwide, our systematic assessment links urban challenges, NbS and outcomes.

Our assessment reveal that specific urban challenges, such as microclimate mitigation and biodiversity loss, were predominantly addressed through specific sets of intuitively related outcomes. Well-being dimensions beyond human health, such as social justice and cohesion and living standards, were predominately reported as secondary outcomes to address other challenges, such as flood risk.

Urban NbS generally yielded positive effects compared to non-NbS. However, the performance varies depending on the considered types of NbS, comparison design types, and temporal scale of the evaluation. Finally, our analysis underscores a notable scarcity of evidence of NbS that report multiple outcomes related to more than one broad category, i.e. health and well-being, regulating ES, or biodiversity.

The identified linkages, as well as the performance reported in the evidence, show a diverse array of urban NbS portfolios, providing supporting information for urban planners, and policymakers to better understand which NbS type could be more pertinent for targeted urban challenges.

## List of Appendix

Appendix A Protocol

Appendix B Study inclusion and exclusion results

Appendix C Supplementary results, figures and tables

Appendix D Systematic map database

## Supporting information

Appendix A Protocol

Appendix B Study inclusion and exclusion results

Appendix C Supplemental results, figures and tables

Appendix D Systematic map database

