## Appendix A Protocol for "Solution to what? Global assessment of nature-based solutions, urban challenges, and outcomes"

### 1 Abstract

The broad framing of the NbS term and the uncertainty about what constitutes an effective NbS hinder its practical adoption (Seddon et al., 2021). The main reason for the vagueness stems from a lack of synthesized evidence on the effectiveness of NbS, entailing the specific problems for which a nature-based solution is needed, and the monitored outcomes (Nature editorial, 2017; Seddon et al., 2020). Therefore, we undertook systematic evidence mapping to consolidate the large, dispersed evidence base on outcomes of NbS for addressing urban challenges.

Our objectives were to collect the data to identify and classify urban NbS and conduct systematic analyses on: 1) relations between urban challenges, NbS types, and their outcomes; 2) the evaluation of the performance of NbS against non-NbS; and 3) assessment of cases reporting multiple outcomes and their interrelationships.

This protocol details the methodology we used to conduct the systematic map, following the Roses (Version 1.0) content guidelines for systematic map protocols (Haddaway et al., 2018). The question scope, search string, and study selection criteria were based on the Context, Intervention, Outcome, Study type (CIOS) framework, which defines the scope of our review without disregarding empirical studies in parts of the world or research fields where the concept is called differently. screened using a stepwise application of inclusion and exclusion criteria at the title/abstract, and full text levels. The screening process was facilitated by using the web tool CADIMA (<https://www.cadima.info>). The extraction of evidence from studies was guided by a coding framework and documented in FileMaker Pro 19. The output of this study is a manuscript synthesizing the systematic map.

### 2 Background

Nature-based solutions (NbS) were defined as “actions to protect, sustainably manage and restore natural and modified ecosystems that address societal challenges effectively and adaptively, simultaneously benefiting people and nature” (IUCN, 2020). We are just at the beginning of systematically analyzing their effects, effectiveness, and provision of co-benefits. Although the implementation of NbS has been increasing dramatically these years, urban planners and managers somehow ignored the identification of the objectives and also lack the understanding of what constitutes successful and sustainable NbS before choosing NbS interventions for implementation.

One reason for this is a lack of synthesis of the evidence on the effectiveness of NbS for urban challenges. To our knowledge, there have been no comprehensive reviews conducted on the effectiveness of urban nature-based solutions for addressing multiple urban challenges based on vast empirical evidence from a global scale.

### 3 Scope, objective, and questions

#### 3.1 Overarching question

What evidence is there for urban nature-based solutions and their outcomes to address urban challenges?

#### 3.2 Secondary questions

- How the NbS is characterized in the urban context?
- What is the relation between urban challenges, NbS types, and outcomes reported in the current evidence?
- What evidence is there showing the comparative performance of NbS?
- What evidence is there reporting multiple outcomes and how are the outcomes related to each other within the multifunctional network?

#### 3.3 Objectives

Our objectives were to collect the data and conduct systematic analyses on:

1) the identification and classification of urban NbS;

2) relations between urban challenges, NbS types, and their outcomes;

3) the evaluation of the performance of NbS against non-NbS;

4) assessment of cases reporting multiple outcomes and their interrelationships.

#### 3.4 Definitions of the question components

We base our protocol on the CIOS framework, adapted from CIMO (Denyer & Tranfield., 2009) and PICOS (James et al., 2016; Moher et al., 2009). This was used to 1) develop a keyword search string, 2) define study selection criteria applied stepwise to filter the search hits, and 3) design a coding framework to delineate what information to extract from the studies.

This framework defines context, intervention, outcome, and study type as the key elements for defining the study scope. The context defines the specific setting in which the intervention will be applied, here urban areas. The intervention itself defines the action to be studied, which here refers to implemented NbS. The outcome frames the range of the effects of the intervention, referring to all the measured and observed effects related to NbS. Finally, the study type defines the study designs of scientific literature, which here is confined to empirical studies.

**Table A1.** Context, Intervention, Outcome, Study type (CIOS) framework used to define key elements for question scope and the inclusion of relevant studies.

| **Context** | **Intervention** | **Comparator** | **Outcome** | **Study type** |
| --- | --- | --- | --- | --- |
| Urban setting | Implemented NbS | N | The effects of NbS interventions | Scientific literature with empirical studies |

### 4. Comprehensiveness of scope for NbS effectiveness

Adapted from the five categories of NbS approaches from Cohen-Shacham et al. (2019), this map includes a list of search terms nesting under the concept of NbS, thus including a broader range of NbS interventions with a broader scope and less bias. Furthermore, this map encompasses a wider range of outcomes related to NbS intervention for both biodiversity and human well-being. Compared with these previous systematic or non-systematic maps, this current systematic review will map a rigorous scientific evidence base with a comprehensive scope of NbS with a broader and innovative search method for retrieving NbS evidence on a global scope.

### 5 Search methods

The systematic map was developed following the RepOrting standards for Systematic Evidence Synthesis (ROSES) for systematic map protocols.

#### 5.1 Scoping process

A scoping process was carried out to inform the search string and study selection criteria.

A scoping exercise was conducted on the Web of Science (WoS) (CORE collection citation indexes) and Scopus databases to build up the search strings. Terms describing the intervention (NbS) were identified and searched iteratively until searches resulted in a suitable number of hits that captured key sources identified from the relevant literature. This stage was processed from August to October 2021.

To increase comprehensiveness, we initially chose both Scopus and WoS. On September 9th, using the previous search strings, 6688 hits were found in WoS, while 7462 were found in Scopus. Both results were exported to CADIMA, after the duplicate removal, 9165 distinct articles were kept in the database.

Using this CADIMA database, we conducted the screening exercise to refine the selection criteria. At the same time, we considered that screening and coding more than 9000 would be unfeasible. During the search training process (August - October), consistently more records were retrieved from Scopus than from WoS, regardless of adding any restrictions or changing search strings. We hence concluded that for our research scope, Scopus has a broader and more up-to-date bibliographic coverage than WoS.

#### 5.2 Search terms selection and comprehensiveness of the search

Our search strings were based on the Context and Intervention elements within CIOS. For Context, search strings composed of terms related to urban areas were applied, such as ‘urban’ and ‘city’. We proposed a broad set of terms captured global scientific research that applied NbS-related thinking without necessarily using NbS terminology.

To cover the breadth of nature-based terms now included under the conceptual umbrella of NbS, the first question to figure out is ‘**What terms can be included under the conceptual umbrella of NbS**’. We identified the search items with the following steps.

1) We adapted the five categories of NbS approaches from Cohen-Shacham et al. (2019). Since it does not include related words about nature and even doesn’t include the keywords ‘nature-based solutions’, **we added one category only for nature-based approaches.**
2) We combined the search terms from current research (Cohen-Shacham et al., 2019; Dick et al., 2020; Nesshöver et al., 2017). To reduce bias, we removed items specific to climate, disaster, risk, flood, and so on. Also, we remove the words on ecosystem types and landscape types such as green roofs and sustainable drainage.
3) Based on the examination of current NbS reviews, terms including nature-inspired solutions, sustainable land management, green spaces, and blue spaces were added. To supplement and remove the words with bias and irrelevant concepts, we reviewed 119 review papers on NbS examining the title, keywords, and abstract to figure out the related concepts that used to represent the same meaning as NbS or as a substitute for NbS. The search for the 119 reviews was conducted in the Web of Science (WoS) database on August 10, 2021, using the search query ‘TS=nature-based solution*’.

After several rounds of scoping exercises and modifications, the search terms used in the final search are shown in the following table.

**Table A2. Categories and specific search** terms used in this study. **Categories adapted from (Cohen-Shacham et al., 2019).**

| **Categories** | **Specific terms used in the search** |
| --- | --- |
| Nature-based approaches | Nature-based solutions, nature-based interventions, nature-inspired solutions, nature-based engineering, nature-based rehabilitation, green solutions |
| Restorative approaches, focus on nature | Ecological restoration, landscape restoration, habitat restoration, land restoration, ecological engineering |
| Issue-specific ecosystem-related approaches | Ecosystem approach, ecosystem-based approach, ecosystem-based adaptation, ecosystem-based mitigation |
| Infrastructure related approaches | Natural infrastructure, ecological infrastructure, green infrastructure, blue-green infrastructure, green space, blue space, green and blue space |
| Management approaches | Sustainable (land) management, best management practice (BMP), natural resource management, ecosystem-based management |
| Ecosystem protection approaches | Area-based conservation approaches, protected area management |

#### **5.3 Search strategy**

We conducted the search in Scopus on 25 October 2021 with search strings. We restricted the search to title content, abstract content, author keywords, and English language. No subject or category refinements were done. Reviews, Proceedings, book chapters, editorials, opinions, commentaries, and perspectives will be excluded from the searches.

**Box A1** Search strings used in Scopus database:

TITLE-ABS-KEY("nature based" OR NBS OR "nature inspired" OR "green solution*" OR "ecosystem approach" OR "ecosystem based approach*" OR "ecosystem based adaptation" OR "ecosystem based mitigation" OR "ecological restoration" OR "landscape restoration" OR "land restoration" OR "habitat restoration" OR "ecological engineering" OR "natural infrastructure" OR "green infrastructure" OR "blue infrastructure" OR "ecological infrastructure" OR "green space" OR "blue space" OR "sustainable management" OR "best management practice" OR BMP OR "ecosystem based management" OR "natural resource management" OR "area based conservation" OR "protected area management") AND TITLE-ABS-KEY(urban OR city OR cities) AND PUBYEAR > 2014

We restricted the published time from 2015 to present for two reasons. First, the scientific research for NbS started in 2015. The concept of NbS and similar concepts are very recent. The scientific community would take time to get more used to these concepts to conduct research on them and use them in the right way. To date, the term NbS has been used mainly in communications targeting policymakers (Cohen-Shacham et al., 2016) and with the exception of one scientific brief (MacKinnon & Hickey, 2009), one proceeding paper (Reguero et al., 2014) and one book chapter(Seidl et al., 2014), it has only recently started to be used in the scientific literature from 2015 (Eggermont et al., 2015; Kabisch et al., 2016; Maes & Jacobs, 2015). Second, there is a need for current recent research on broader NbS. After NbS entered the scientific sphere, it is currently transferring, transforming, and working on and with ideas, concepts, and methods developed previously (Hanson et al., 2020). Emerging of the NbS symbolizes the outset of the development of more sustainable and inclusive cities, balancing anthropocentric and ecocentric values (Randrup et al., 2020). Older green terms, for instance, green space change the research focus. We hence limited the temporal extent of the search to ensure that only research that is currently relevant and closer to NbS is included in the analysis.

Citation outputs from Scopus were exported to the searched records by relevance yearly and then exported to CADIMA for removing duplicates and screening.

### 6 Article screening study inclusion criteria

#### 6.1 Screening strategy

Selection criteria were applied stepwise during screening in CADIMA, first screening citation titles and abstracts, followed by full texts. Before screening, we progressively refined the criteria to ensure they were clear to all reviewers and interpreted consistently. Selection criteria were refined during the abstract screening and full-text assessment for coding.

A series of eligibility and exclusion decisions regarding CIOS (Table A2) was consistently applied to determine the relevance of articles, focusing on the study context, a defined intervention, study outcome, and study type.

We carried out single reviewer screening cautiously by checking any uncertain decisions with the other co-authors to reach an agreement throughout the process. Decisions at each stage of screening were conservative, by assessing studies for which inclusion eligibility was unclear.

To improve the clarity of the criteria, we conducted two tests for **study selection consistency** with 30 papers among 4 reviewers in September 2021 and 50 papers among 3 reviewers in October 2021, respectively. The exercises were conducted based on 9165 records including records from WoS (6688) and Scopus (7462) on 9 September 2021, and we used the consistency check function in CADIMA. A total of 80 articles were randomly selected using CADIMA. In the first round, 30 articles were selected, and in the second round, an additional 50 articles were chosen. The decision could be made by three choices: Yes, unclear, or No. The decision was made first by the four categories including context, intervention, outcome, and study type, then followed by the final decision of this article. The working files of consistency checks were downloaded from CADIMA for all of us to work on and the results were summarized (shown in Fig. A1).

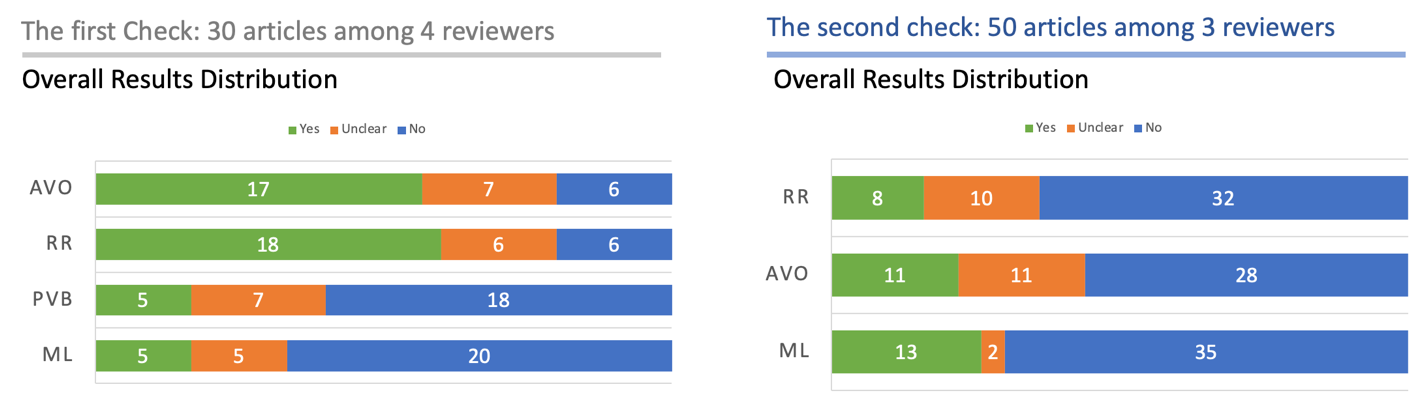

**Fig. A1** Summary of consistency check results.

We reviewed any emerging, conflicting selections based on each criterion and revised the screening strategy and selection criteria for clarity. As shown in Fig. A1, the general consistency of the second check was improved a lot compared with the first one. After these tests, the main refinement involved defining the NbS intervention more precisely and excluding publications that do not study specific interventions, such as generic green spaces. Another main decision was to exclude publications that did not yield primary data in evaluation.

Considering the broadness of interventions and the variety of evaluation methodologies applied in the studies, we conducted a substage of full-text screening with a detailed examination of the evaluation methods for eligibility for coding. This involved a preliminary coding process of key elements such as NbS interventions and outcomes with applied indicators. For work feasibility, we selected 75% of the 691 articles (521 articles) that were most relevant to our study scope. As described in Section 5.3, the article records ranked by relevance in Scopus were exported separately for each publication year (see distribution of selected publications below). To extract the most pertinent records for our study scope, we selected the top 75% of records based on each publication year. Following these steps, our selection of these 521 articles provided the best representation of the literature suitable for our research topic. During the process, we refined the criteria based on our continuous discussions on the interpretation of the scope of NbS, i.e., what is and what is not an NbS. In addition, a specific check on the evaluation methods was conducted, including the precision of data, measurements, indicators and key results. These led to the refinement of outcome criteria as listed in Table A3. As a sequence, we finally refined the criteria (Table A3), and a further 160 publications were removed for coding and evidence mapping from 521 processed articles. The detailed explanations for exclusion are provided in Appendix C.

**Table A3.** Distribution of selected studies in each publication year in 75% of total included papers after full-text screening.

| **Publication year** | **Total number** | **75% processed** |
| --- | --- | --- |
| 2022 | 2 | 2 |
| 2021 | 146 | 111 |
| 2020 | 132 | 99 |
| 2019 | 112 | 84 |
| 2018 | 96 | 72 |
| 2017 | 89 | 67 |
| 2016 | 70 | 53 |
| 2015 | 44 | 33 |
| **In total** | **691** | 521 |

**Table A4.** Inclusion and exclusion criteria used in the review, based on the Context, Intervention, Outcome, and Study type (CIOS) framework.

| **Eligibility criteria** | **Inclusion** | **Exclusion** |
| --- | --- | --- |
| **Context (Does the study deal with challenges in urban areas? )** | involve outdoor landscape in the urban context | not involve the urban context, e.g. regional, national, continental processes and indoor plants, but experiments with the urban setting are included |
| **Intervention (Does the study involve any NbS intervention?)** | Intervention in relation to nature-based solutions includes two general types:  1) The presence of **specific types of spatial units** ( e.g. urban forests, urban parks, urban trees, green walls). For example, the creation of green roofs and walls to mitigate rainwater runoff. 2) **The actions to apply measures or techniques to alter the spatial units.** Examples include developing structural attributes/microscale elements of ecosystems such as enhancing tree density and vegetation species to improve biodiversity and cultural experiences. | 1) Interventions not employing nature (without the presence of biotic components). For instance, “porous asphalt”, replicates the natural process of water infiltration, yet it does not necessitate the presence of urban nature. Engineered solutions, such as large dams and inter-basin transfers through pipes.  2) Investigating general green spaces (e.g. green areas, stormwater management systems) or general land use types (e.g. forests and shrubs) without involving any links to specific actions or mentioning any specific spatial unit types (e.g., urban forest, parks, street trees). Examples include investigating the spatial distribution of urban green space changes (increase or reduction) using general land use data to indicate outcomes for land surface temperature and rainwater runoff reduction. |
| **Outcome (Does the study directly assess the effects in relation to urban nature?)** | Studies involve empirical evidence (i.e., studies yielding primary data) with a quantitative or qualitative evaluation of implemented NbS intervention.  Notes: This includes any outcomes provided by NbS, that might be related to biodiversity gains or human-wellbeing benefits. | Studies not undertaking any direct assessment of NbS intervention's impact on human well-being or biodiversity. Including 1) Studies not aiming to provide an assessment of NbS effectiveness. Examples include,   where data was only used to characterize green spaces/ecosystem services types (Atasoy et al., 2018) and map the distribution.  where data was used to investigate motives to use parks/use patterns/ preferences of parks and the influencing factors.  where the main aim is to investigate confounding factors(including NbS characteristics such as age, area, tree coverage percentage, etc. ) that have impacts on the outcomes.  2) Studies investigating **confounding factors,** which precluded firm conclusions on the outcomes in relation to NbS intervention. Examples include where recorded bird biodiversity degradation could have resulted either from plant diversity or from noises from urban settings. 3) Studies focusing on **mechanism research /process research,** thus having no direct clue for the effectiveness of NbS intervention (need to indicate). Examples include the investigation into the impact of individual leaf traits on traffic-generated PM accumulation. |
| **Study type（Does the study deal with empirical studies involving quantitative or qualitative evaluation? ）** | Studies involve empirical evidence with quantitative, or qualitative evaluation on **implemented NbS intervention**. We defined empirical studies as those reporting direct observations or measurements of the effects of an intervention (i.e., studies yielding **primary data**). | 1) Studies on effectiveness **scenarios/forecasts modeling**, unless where pre-intervention reference conditions were simulated using a model, but where the local data of post-intervention were employed.  2) Studies on **planning approaches** or tools to support the implementation of NbS (e.g. prioritizing which interventions to implement and where) or to preserve/ monitor urban ecosystems (this also will be excluded with outcome criteria, and maybe with intervention criteria), 3) Studies proposing **new conceptual frameworks, methodology, or developing tools for evaluation** (Zhang et al., 2020), unless they also provide relevant empirical evidence (e.g. using empirical studies to demonstrate new methodologies). (this also will be excluded with outcome criteria, and also with intervention criteria) |

#### 6.2 Transparency/Reporting

We used the ROSES flow diagram (N. Haddaway et al., 2017) to record the flow of citations through searching, screening, coding, and synthesis. A list of studies excluded at the full-text level, and the reasons for their exclusion, is available. Our evidence map is accessible on CADIMA, by searching for the title of this research at <https://www.cadima.info/index.php/area/evidenceSynthesisDatabase>. The screening process outcomes were available there. In addition, we provided a list of articles excluded during data extraction and coding with explanations of why they could not be processed.

### 7 Data extraction and coding

#### **7.1 Coding strategy**

The extraction and coding of evidence from articles were guided by a coding framework with definitions (see full details in Appendix D). We defined a list of coding definitions to standardize coding decisions and improve consistency among those involved in coding articles. We proposed a list of definitions regarding NbS types, urban challenge categories, and outcome domains to categorize the database and also defined the performance assessment by comparators and temporal scale. We iteratively modified the framework and coding definitions by applying the framework to a set of 26 benchmark studies (before April 2021). This framework is a broad-level, overarching framework designed to have sufficient flexibility to capture the heterogeneous evidence base on the outcomes of nature-based interventions for addressing urban challenges.

The data extraction was conducted by two researchers, ML and JF. JF was trained in coding with the predefined coding definitions. The coding of 97 out of 361 publications were cross-checked by two coding contributors. Any confusions were addressed and checked for consistency among the whole research team. In the final stage, ML validated all the work from the JF and compiled the coding database.

#### **7.2 Coding framework**

We designed a coding framework to extracte the following data: 1) Article Bibliographic information, 2)Basic information about the intervention, including locations, NbS types and urban challenges, and multifunctionality, 3) Outcome evidence, including outcomes reported and performance compared to non-NbS. The whole framework was a provided in Appendix D Systematic map database.

In general, each unique combination of intervention type, and location (urban setting) in a study were recorded as a separate intervention case. Each article would have one or more intervention cases, each having one or more possible outcomes relating to an intervention case.

Each article was assigned a unique Article ID, a continuous number based on the CADIMA ID. The numbering follows an order where it first increases in descending order of publication year and then in descending order of Scopus relevance.

NbS case entry. We assigned each intervention (cases) a unique Study ID. When there are multiple types of NbS interventions in one article, 1) If the evaluation aims to be done across the types, record all of them as separate NbS cases; 2) If the evaluation aims for one type and only uses the other types as control comparators, only record the initial NbS type as a case study and record other types in the box of comparison assessment with other types. When there are gradient management measures of green spaces: 1) If the evaluation aims to be done across the gradient of management measures, record extreme conditions as two types of NbS. 2)If the evaluation aims for one type, record only the other type as control comparators. For example, research comparing along the gradient of the management measures covering different mowing frequencies (intensive-normal- extensive) of urban lawn, is indeed comparing across different types of NbS. In that case, we will record extreme conditions as two types of NbS, which are intensive management and extensive management, as separate cases and the comparison will be ‘Other type’.

Outcome entry. For each case study, we extracted all outcomes evaluated by indicators that were presented in the results section of the publications. The results of input data relating to the interventions were recorded as outcomes. We only recorded quantified outcome assessments, thus those using general words without numeric description were included. Each outcome might be evaluated through multiple variables.

#### **7.3 Coding classifications and definitions**

##### 7.3.1 Nature-based solutions classification

Coding for urban NbS types followed our definitions of the classification of urban NbS (See Table D1 below). Each of the NbS identified in the database was coded based on the proposed classification as depicted in Fig.3 in the article.

For defining the classification of urban NbS, we extracted the description of the intervention as specified by the authors. Make the description as specific as possible to allow more fine-grained categorization of intervention types after initial data extraction.

The classification was established by a deductive coding based on the extracted data of NbS characteristics of 547 cases. To enable evaluating the linkage to urban challenges for this research, our classification of urban NbS was only based on the physical features (e.g., forms of vegetation), without considering other aspects such as the purposes of the actions. Note that original articles might use other names, but we only rely on the descriptions of the features and categorize them with our classification. Thus, each type represents cases that have been associated with various frequently used names in research. For example, under the semi-natural park type, one can find cases using more specific and unique names such as public cemetery, pocket park, and theme park (see Frequently used names, in Table A5).

The classification first build on the two categories proposed by Castellar et al. (2021). NbS units are standalone green technologies or green urban spaces and NbS interventions are interventions in existing ecosystems and NbS units. Our classification adapts the categories based on the physical features of NbS, regardless of purpose and other considerations. We proposed the list of types based on our examination on the physical features depicted in the studies. See definitions in Table A5.

**Table A5.** NbS classification with descriptions and frequently used names under each categories.

| NbS Categories | **Featured groups** | **Description** | **NbS Unit Type** | **Description** | **Frequently used names by authors** |
| --- | --- | --- | --- | --- | --- |
| NbS Units | **Vegetation** | Vegetation NbS Units refers to a group of unit types dominated by vegetation feature. The types are differentiated by the specific forms of vegetation. | **forest & tree** | vegetation based, dominated by trees | forest, urban trees |
|  |  |  | **grassland** | vegetation based, dominated by grass | grassland & meadow, lawn, sport field |
|  |  |  | **farmland** | vegetation based, dominated by crop | farm |
|  |  |  | **shrubland** | vegetation based, dominated by shrub | hedge |
|  |  |  | **Mixed vegetation** | vegetation based, mixed types | revegetated sites |
|  | **Water** | Water NbS Units refer to a group of types dominated by water features. The types are differentiated by forms of water regarding the morphology of water and the presence of vegetation. | **river & canal** | water based, featured by linear water area such as rivers and canal | river, canal |
|  |  |  | **naturalized wetland** | water based, featured by natural wetland with vegetation | naturalized wetland |
|  |  |  | **stormwater basin** | water based, semi-natural, temporal existence of water, landscaped depressions linked to engineered facilities(i.e., roads and drainage system) | swale, retention/detention cells, rain garden |
|  |  |  | **constructed wetland** | water based, constructed units, linked to engineered facilities(i.e., roads and drainage system) | constructed wetland |
|  | **Hybrid** | Hybrid NbS Units refer to a group of types with a hybrid feature of vegetation, water, and engineered features. The types are differentiated by the proportion of the 3 features present in the units. | **natural park** | hybrid with built area, mixed vegetation types, sometimes with water features, open area, medium coverage of grey area, featured by nature, only have low percentage of grey coverage | wetland park, forest park |
|  |  |  | **general park** | hybrid with built area, mixed vegetation types, sometimes with water features, open area, medium coverage of grey area, can not tell if it's dominated by natural features or if more recreational facilities are included. | park |
|  |  |  | **semi-natural park** | hybrid with built area, mixed vegetation types, sometimes with water features, open area, medium coverage of grey area, have medium to high coverage of grey area | pocket park, theme park, neighborhood park, public cemetery |
|  |  |  | **garden** | hybrid with built area, mixed vegetation types, dominated by lawn, flowerbed, crops, sometimes trees and water features, often enclosed area, medium coverage of grey area | garden, domestic garden, community garden, botanic garden, cultural garden |
|  | **Engineered** | Engineered NbS Units refer to a group of types supported by engineered features. The types are differentiated by specific forms of engineered components. | **green alley & roadside green** | street based, mixed vegetation | green alley, roadside plantation, street trees, attached green spaces |
|  |  |  | **green square** | square based, mixed vegetation types, sometimes water features facilities, open area, higher coverage of grey area | green square |
|  |  |  | **green roof & wall** | building based, combined with building structure, mixed vegetation types, facilities | green roof, green wall, green structure |
|  | NA | Others representing the cases did not specify the components of features. | **Others** | The description of the NbS characteristics is general(only based on the intention to use or can refer multiple features)and difficult to identify based on our criteria. | green corridor, reserve, brownfield |
| NbS Categories | **Featured group** | **Description** | **NbS Measure Type** | **Description** | **Frequently used names by authors** |
| NbS Measures | Plant | Plant (Abiotic) measures include a diverse set of techniques to modify the abiotic components | **Mowing/cutting** | The act of leveling or cutting down grass, grain, and other etc., with a mowing machine or scythe | changing mowing frequency of lawns, cutting down the hedge along a pathway |
|  |  |  | **Revegetation/planting** | The process of replanting and rebuilding the soil of disturbed land. | revegetation/planting on the riverbank, forest, abandoned area, introduction of plants (green algae), Perch treatments |
|  |  |  | **Managing native/invasive plants** | Management of invasive and local plant species. | replacing exotic plant species with wild/local species, removal of woody invasive species and planting with native tree seedlings, introduction of indigenous tree species |
|  |  |  | **Macrophyte harvesting** | Management of macrophyte biomass via harvesting for maintaining eutrophic inland waters | macrophyte harvesting |
|  | Soil | Soil measures include a set of soil bioengineers to modify the features of soil | **Mulching** | The process of covering the open surface of the ground by a layer of some external material | mulching |
|  |  |  | **Soil rehabilitation** | The process of carrying out a surface soil tillage, laying a new surface layer of fertile topsoil, and sowing grass species on the new surface | Soil rehabilitation, Soil decompaction and compost amendment, compost, soil decompaction and compost amendment |
|  | Abiotic | Abiotic measures include a set of measures applied in abiotic components such as water and engineered infrastructure. | **Creation of bypass channels** | Creation of inner facilities of river channel | construction of a bypass channel |
|  | NA | Combination of abiotic, plant and soil measures | **Combined measures** | Combination of multiple specific measures defined above | River or forest combined restoration intervention |

##### 7.3.2 Urban challenge categories

To propose the classification, we first recorded the description of the author (used to define the categories). To get this information, one needs to check the background and the introduction of the project. Record a UC if the authors state that it is **an existing issue**. The intervention outcomes might match the UC or not. (e.g. 100% match: lack of habitat for bird biodiversity is an issue in the area and the study measures the abundance and richness of birds).

We adapted the typology of 23 urban challenges from existing categories (Babí Almenar et al., 2021; Dumitru & Wendling, 2021).To explore the multifunction and potential disconnected from specific challenges, we also include ‘multiple urban challenges’ and ‘no specific challenges’ in the categories:

a. Health and well-being – Issues related to physical and mental health, general subjective well-being, or general health and well-being.
b. Biodiversity loss – Issues related to the loss of biodiversity species. Also coded as biodiversity loss when the author explicitly identified the issue as biodiversity loss.
c. Urban heat island – Issues related to heat waves and urban heat islands.
d. Habitat degradation – Issues related to the loss of ecosystem types (i.e., forest, floodplain, wetland) because of urban expansion and construction.
e. Water pollution and shortage
f. Greenhouse gas emission – Issues related to global warming and anthropogenic CO2 emissions.
g. Flood risk
h. Soil degradation
i. Air pollution
j. Lack of open green spaces - Issues related to the lack of green space and scarcity of common areas caused by urban expansion.
k. Pollinator and pollination loss
l. Food insecurity
m. Urban shrink and decline – Issues related to growing vacant land/brownfields accomplished by urban decline.
n. Public perception and participation – Example includes the social acceptance of urban wastewater recharge projects.
o. Urban development issue
p. Ecosystem disservices
q. Social justice and cohesion issue
r. Natural hazards – Examples include tropical cyclones and dust storms in our database.
s. Noise pollution – Issues mainly related to traffic noise.
t. Cultural heritage protection
u. Climate change
v. Multiple urban challenges - More than one urban challenge was stated.
w. No specific challenges – When the study does not explicitly state any urban issues.

##### 7.3.3 Outcome domains

We detailed the evaluated outcomes related to the intervention as denoted by the author.

The outcome should be used to show the effects or impacts on social challenges surrounding biodiversity and human well-being. Only the outcomes evaluated using variables or proxies are recorded in our database.

To identify a list of outcome domains to represent ES, human well-being benefits, and biodiversity gains, we applied a bottom-up method based on outcome domains described by the authors and the indicators used, see Table D1. The description of well-being domains was modified based on Lisa M. Smith et al. (2013), and Dick et al. (2020).

Outcome domains are mainly based on the evaluation aim. Note that the same indicator can be used for various outcomes. For example, physiological indicators such as blood pressure can be used for ‘physical health’ assessment when the research evaluates the impact on cardiovascular function and hypertension. They also can be used to evaluate the impacts on anxiety or nerve activity, which refers to ‘mental health’ linked to stress. If using indicators related to biodiversity measurements, but aiming for evaluating water quality improvement, code as water quality enhancement. Only when the author doesn’t have a clearly defined outcome domain, will we code based on the indicators.

**Table A6** The identified outcome assessment domains with definitions and frequently used indicators.

| **Broad category** | **Domain** | **Definition(scope)** | **Frequently used/typical indicators used for the assessment** |
| --- | --- | --- | --- |
| Biodiversity | Plant biodiversity | Conservation of plant community | Species richness and diversity of plants such as trees and grass |
|  | Bird biodiversity | Conservation of bird community | Species richness and diversity of birds |
|  | Invertebrate biodiversity | Conservation of invertebrate community | Species richness and diversity of invertebrate communities, such as insects and arachnids |
|  | Mammal biodiversity | Conservation of mammal community | Species richness and diversity of mammals |
|  | Fish biodiversity | Conservation of fish community | Species richness and diversity of fishes |
|  | Undefined biodiversity | Conservation of generic biodiversity | Biota richness and perceived biodiversity |
| Health and well-being | Mental health | Psychological, emotional, and mental health | Improved mood status, cognitive development, stress recovery, and reduced depression risk |
|  | Physical health | Physical health | Reduction of disease occurrences, mortality rate, and improvement in maternal health, child health, body status |
|  | General health | Physical and mental health | Reported general health conditions |
|  | Physical activities | Improved level of outdoor active activities such as cycling and walking | The probability, the distance, the time of activities |
|  | Recreation | Leisure activities or enjoyment of natural scenery | Accessibility, facility supply, duration/frequency of activities, and perceived benefits |
|  | Aesthetic appreciation | Aesthetically valuable experiences derived from the attractiveness | Perceived aesthetic value |
|  | Cultural and spiritual experience | Cultural, societal, and traditional values of natural resources | The reported value of the sense of home, connection to nature, cultural identity and heritage, spiritual or religious beliefs |
|  | Life satisfaction and happiness | Contentment with our life | Self-reported happiness and satisfaction with life |
|  | Social justice and social cohesion | Interactions between individuals, within and between groups | The level of conflict mitigation, relationships, connectedness, ability to work together, ability to help others, and trust |
|  | Security and safety | Personal safety, resource security, and human rights, both perceived and actual | Reported sense of security and crime rate |
|  | Living standards | Economic benefits for the well-being of individuals and society | Status of income, poverty, employment, wealth, savings, payments |
|  | Education | Benefits from formal and informal knowledge transfer | Quality of education, degrees awarded, livelihood skills, and environmental knowledge |
|  | Overall well-being | Overall subjective well-being | Reported well-being includes more than one scope of other well-being categories such as mental health, mood status, and life satisfaction. |
| Regulating ES | Microclimate regulation | Regulation of local climate to reduce the risk of heat stress and other microclimate risks | Level of temperature, humidity, ventilation, and transpiration |
|  | Greenhouse gas mitigation | Climate change regulation by improving carbon sequestration or mitigating GHG emissions | Carbon storage and sequestration |
|  | Flood mitigation | Ecosystems attenuate stormwater flooding as rainfall is intercepted and stored to reduce the flooding risks | Peak flow rate and runoff volume |
|  | Water quality enhancement | Regulation of the physical-chemical condition of freshwaters, to improve water quality | Aquatic habitat condition and the physical and chemical characteristics of freshwater |
|  | Air purification | Reduction of health risks associated with poor air quality | The concentration of air pollutants such as PM10, PM2.5, and NO2 |
|  | Noise abatement | Mediation of nuisances of noise, to improve the well-being of living systems | Noise test by a decibel meter |
|  | Soil quality enhancement | Regulation of physical, chemical, and biological conditions of soil to improve soil quality that can ensure people’s usage | Soil compaction and soil biomass |
|  | Pollination | Ecological function supporting food production and maintaining flowering plant diversity | Pollinator diversity and visiting frequency, and amount of pollinator habitat |
|  | Nature-borne disease mitigation | Mitigation of the increased risk of disease from exposure to nature | The potential risk of exposure to ticks, mosquitoes, and pollens |
| Others | Resource efficiency | Cost reduction for installing and maintaining NbS | Monetary and material consumption such as water consumption volume, life cycle cost, financial/economic cost |
|  | Food production | Food supply derived from edible green spaces | Fruit and vegetable productivity |

##### 7.3.4 Performance

1) Comparisons

**The performance is assessed based on the effects using non-NbS comparators, i.e., comparison with no NbS intervention(e.g. green roof versus a grey roof, or before versus after a walk in an urban park).** This group includes the comparison between pre-and post-intervention, and post-intervention compared to reference sites representing pre-implementation conditions, post-intervention compared to scenarios based on computer simulation without intervention, and comparison between with and without a post-NbS intervention from a stakeholder perspective.

If outcome assessments used other NbS types as comparators (e.g. a community park against a national park, or urban lawns mowed intensively against extensively)., we marked them for further exploration.

Studies involve no comparison was also marked. This group includes measuring performance against the desired threshold (e.g. such as water quality standard) and natural references(e.g. natural rivers or forests in the countryside). In this circumstance, it's hard to decide if the outcome is positive or negative when the desired threshold is not met or not compared with the natural references.

The comparison between NbS and non-NbS controls were further defined and classified. By further examining the reported effects, we extracted information on the detailed experimental or study design of the compared research and clustered them into 4 distinct groups:

a. **Control site**: comparing different spaces with and without NbS intervention, such as built areas and bare roofs;
b. **Before and after**: comparing conditions before and after implementation of NbS or engaging with the NbS;
c. **Green exposure**, comparing conditions with high and low exposure to NbS, encompassing various factors related to the presence and influence of greenery in an individual’s surroundings which include accessibility, duration of time spent in green spaces, and frequency of exposure.
d. **Simulated non-NbS:** comparing post-intervention outcomes with simulated scenarios without intervention, for instance, a scenario with all vegetation removed.
e. **Combination:** In cases where the comparator design for a specific outcome incorporates more than one of the defined comparator design types mentioned above, it is categorized as a combination.

2) The performance against non-nature-based solutions comparators

The performance was assessed by reported effects (positive, negative, neutral, mixed, or unclear outcomes). The effects were coded based on the following definitions, including the conditions where more than one indicator is used for one aiming outcome domain:

a. Positive

When a positive result is indicated, i.e. the climate change impact is reduced. Code as such if both positive and neutral effects are reported. However, if both positive and neutral effects are reported over the temporal scale, code as mixed.

b. Negative

When a negative/adverse result is indicated, i.e. the climate impact is increased/worsened. Code as such if both negative and neutral effects are reported. However, if both negative and neutral effects are reported over the temporal scale, code as mixed.

c. Neutral

When neither positive nor negative result is reported.

d. Mixed

When both positive and negative results are reported. Results can vary based on location variations or across time. If some results are positive (or negative) and others are unclear, also code as ‘mixed.’ OR when comparing the outcome with multiple types of NbS one type ranks in the middle or has a lower ranking compared to the best one. OR the

e. NA

When the authors do not derive an explicit conclusion through any comparison.

3) The temporal scale of performance

Furthermore, we meticulously documented the **temporal scale** associated with these effects, discerning between two key categories:

a. Cross-sectional effect: The effect of NbS is measured and comparatively evaluated at a specific single time point (excluding measurements for a before-implementation control), providing a snapshot of the effect within a specific and delimited period. The study design involves one data collection wave and applies a single cross-sectional analysis.
b. Effect over time: The effect of NbS is assessed based on multiple measurements and comparative evaluation of the outcomes across various time points, capturing changes in performance over a designated temporal period. The study design involves two or more data collection waves and applies either repetitive cross-sectional analysis or longitudinal observation.

Please note that an effect evaluated by a before-and-after comparison (involving two single time points) should be classified as cross-sectional unless it measures ‘after’ outcomes multiple times considering the dynamic changes or process between the reported outcomes.

The performance involving multiple effects over time coded as positive shows to be increasingly or consistently effective in addressing certain challenges over time, for example, increasing biodiversity enhancement with time. Those coded as mixed means that both negative and positive effects were reported within the assessments for various time points.

#### **7.4 Supplementary coding for the selection of 75% most relevant studies**

To address the concerns regarding the potential impact of the excluded 25% on current reported findings, we appraised these 170 articles, by screening titles, abstracts, and full texts and preliminary coding. Based on the quick examination, we found that 62% of the 170 were potentially qualified for further coding and synthesis, a lower inclusion rate compared to the 69% observed in the most relevant 75% (as depicted in Figure 1 in the manuscript, based on the processing of 521 articles). This reaffirmed the efficacy of our relevance-based selection.

For these potentially qualified papers, we roughly extracted the information related to the geographic location, urban challenges, NbS, and outcomes and coded with our defined coding categories (Table 4). We compared the new data by adding the basic coding results, which present the potential full data of the database. As shown in Tables 5A-D, even though subtle differences were observed, they did not reverse our current findings on the most reported and under-representative categories, such as geographical distribution.

**Table A7 Coding results based on 103 articles potentially qualified for NbS case coding.** Note that the calculation is based on the number of articles rather than separate cases.

**A) Geographical distribution-Continent**

| **Continent** | **Number of articles** |
| --- | --- |
| Asia | 35 |
| Europe | 35 |
| North America | 24 |
| Africa | 4 |
| Oceania | 4 |
| South America | 1 |
| Grand Total | 103 |

B) Urban challenges

| **Urban challenge categories** | **Number** |
| --- | --- |
| No specific challenges | 22 |
| Health and well-being | 21 |
| Urban heat island | 11 |
| Water pollution and shortage | 11 |
| Habitat degradation | 9 |
| Flood risk | 7 |
| Biodiversity loss | 7 |
| Air pollution | 5 |
| Greenhouse gas emission | 3 |
| Lack of open green spaces | 2 |
| Pollinator and pollination loss | 2 |
| Social justice and cohesion issue | 1 |
| Natural hazards | 1 |
| Multiple challenges | 1 |
| Grand Total | 103 |

C) Nature-based solutions unit types

| **NbS unit types** | **Number** |
| --- | --- |
| Forest & trees | 21 |
| Green roof & wall | 14 |
| Garden | 11 |
| General park | 10 |
| Stormwater basin | 10 |
| River & canal | 9 |
| Grassland | 7 |
| Constructed wetland | 7 |
| Green alley & roadside green | 6 |
| Naturalized wetland | 5 |
| Natural park | 4 |
| Others | 3 |
| Semi-natural park | 1 |
| Grand Total | 108 |

D) Outcome domains

| **Outcome domains** | **Number** |
| --- | --- |
| Water quality enhancement | 23 |
| Microclimate regulation | 14 |
| Mental health | 13 |
| Plant biodiversity | 9 |
| Invertebrate biodiversity | 7 |
| Overall well-being | 7 |
| Air purification | 6 |
| Flood mitigation | 6 |
| Greenhouse gas mitigation | 5 |
| Physical health | 5 |
| Physical activities | 4 |
| Cultural and spiritual experience | 3 |
| Soil quality enhancement | 3 |
| Social justice and social cohesion | 2 |
| Resources efficiency | 2 |
| Aesthetic appreciation | 2 |
| Recreation | 2 |
| Pollination | 2 |
| General health | 1 |
| Living standards | 1 |
| Mammal biodiversity | 1 |
| Security and safety | 1 |
| Food production | 1 |
| Education | 1 |
| Grand Total | 121 |

**Table A8** Results of the comparison to the current database.

A) Continent

| **Continent** | **Number** | **Percentage** | **New Number** | **Percentage** |
| --- | --- | --- | --- | --- |
| Asia | 181 | 33% | 216 | 28% |
| Europe | 167 | 31% | 202 | 26% |
| North America | 115 | 21% | 139 | 18% |
| Oceania | 39 | 7% | 43 | 6% |
| South America | 28 | 5% | 29 | 4% |
| Africa | 16 | 3% | 20 | 2% |
| Grand Total | 546 | 100% | 649 | 84% |

B) Urban challenges

| **Urban challenge Categories** | **Number** | **Percentage** | **New Number** | **Percentage** |
| --- | --- | --- | --- | --- |
| Health and well-being | 140 | 26% | 161 | 25% |
| Biodiversity loss | 103 | 19% | 110 | 17% |
| Urban heat island | 68 | 12% | 79 | 12% |
| No specific challenges | 40 | 7% | 62 | 10% |
| Habitat degradation | 33 | 6% | 42 | 6% |
| Multiple | 25 | 5% | 26 | 4% |
| Water pollution and shortage | 20 | 4% | 31 | 5% |
| Greenhouse gas emission | 19 | 3% | 22 | 3% |
| Flood risk | 18 | 3% | 25 | 4% |
| Soil degradation | 17 | 3% | 17 | 3% |
| Air pollution | 16 | 3% | 21 | 3% |
| Lack of open green spaces | 13 | 2% | 15 | 2% |
| Pollinator and pollination loss | 10 | 2% | 12 | 2% |
| Food insecurity | 5 | 1% | 5 | 1% |
| Urban shrink and decline | 4 | 1% | 4 | 1% |
| Public perception and participation | 3 | 1% | 3 | 0% |
| Urban development issue | 3 | 1% | 3 | 0% |
| Ecosystem disservices | 3 | 1% | 3 | 0% |
| Social justice and cohesion issue | 2 | 0% | 3 | 0% |
| Natural hazards | 2 | 0% | 3 | 0% |
| Noise pollution | 1 | 0% | 1 | 0% |
| Cultural heritage protection | 1 | 0% | 1 | 0% |
| Climate change | 1 | 0% | 1 | 0% |
| Grand Total | 547 | 100% | 650 | 100% |

C) Nature-based solutions unit types

| **NbS unit types** | **Number** | **Percentage** | **New Number** | **Percentage** |
| --- | --- | --- | --- | --- |
| Forest & trees | 86 | 16% | 107 | 16% |
| Semi-natural park | 72 | 13% | 73 | 11% |
| General park | 69 | 13% | 79 | 12% |
| Garden | 58 | 11% | 69 | 11% |
| Grassland | 52 | 10% | 59 | 9% |
| Green roof & wall | 49 | 9% | 63 | 10% |
| Green alley & roadside green | 37 | 7% | 43 | 7% |
| Stormwater basin | 29 | 5% | 39 | 6% |
| Natural park | 26 | 5% | 30 | 5% |
| River & canal | 14 | 3% | 23 | 4% |
| Naturalized wetland | 11 | 2% | 16 | 2% |
| Others | 11 | 2% | 14 | 2% |
| Farmland | 10 | 2% | 10 | 2% |
| Green square | 8 | 1% | 8 | 1% |
| Constructed wetland | 7 | 1% | 14 | 2% |
| Mixed vegetation | 5 | 1% | 5 | 1% |
| Shrubland | 3 | 1% | 3 | 0% |
| Grand Total | 547 | 100% | 655 | 100% |

D) Outcome domains

| **Outcome domains** | **Number** | **Percentage** | **New Number** | **Percentage** |
| --- | --- | --- | --- | --- |
| Microclimate regulation | 94 | 12% | 108 | 12% |
| Mental health | 70 | 9% | 83 | 9% |
| Plant biodiversity | 69 | 9% | 78 | 8% |
| Flood mitigation | 49 | 6% | 55 | 6% |
| Soil quality enhancement | 47 | 6% | 50 | 5% |
| Invertebrate biodiversity | 45 | 6% | 52 | 6% |
| Greenhouse gas mitigation | 43 | 5% | 48 | 5% |
| Water quality enhancement | 38 | 5% | 61 | 7% |
| Bird biodiversity | 35 | 4% | 35 | 4% |
| Air purification | 34 | 4% | 40 | 4% |
| Physical health | 34 | 4% | 39 | 4% |
| Pollination | 26 | 3% | 28 | 3% |
| Recreation | 25 | 3% | 27 | 3% |
| Living standards | 22 | 3% | 23 | 3% |
| Cultural and spiritual experience | 19 | 2% | 22 | 2% |
| Resource efficiency | 19 | 2% | 21 | 2% |
| Social justice and social cohesion | 18 | 2% | 20 | 2% |
| Nature-borne disease mitigation | 18 | 2% | 18 | 2% |
| Overall well-being | 12 | 2% | 19 | 2% |
| General health | 12 | 2% | 13 | 1% |
| Aesthetic appreciation | 12 | 2% | 14 | 2% |
| Food production | 10 | 1% | 11 | 1% |
| Physical activities | 9 | 1% | 13 | 1% |
| Education | 8 | 1% | 9 | 1% |
| Life satisfaction and happiness | 8 | 1% | 8 | 1% |
| Mammal biodiversity | 7 | 1% | 8 | 1% |
| Security and safety | 6 | 1% | 7 | 1% |
| Noise abatement | 4 | 1% | 4 | 0% |
| Undefined biodiversity | 4 | 1% | 4 | 0% |
| Fish biodiversity | 2 | 0% | 2 | 0% |
| Total of the outcomes | 799 | 100% | 920 | 100% |

### 8 Data mapping and synthesis

The evidence was characterized through descriptive numbers at two levels: case level and outcome level. The urban challenges, location, and NbS types were counted on the case level while the outcomes and performance assessments were counted on the outcome level. Two-dimensional evidence matrices make explicit linkages between urban challenges, NbS, and outcomes. Information on the performance assessment permitted characterization of the proportion of studies by comparator types and the effects by Non-NbS and Other NbS comparators.
