## Appendix C Supplemental results, figures and tables for "Solution to what? Global assessment of nature-based solutions, urban challenges, and outcomes"

### 1 Supplemental results

1.1 NbS measures and their combinition with units

The most frequently identified measures are combined measures (18), followed by mowing/cutting (16), and revegetation/planting (13) (Fig. S1A). The most reported combination of measures and units is mowing/cutting in grasslands, followed by combined measures in rivers and revegetation/planting in forests (Fig. S1B).

A)	B)

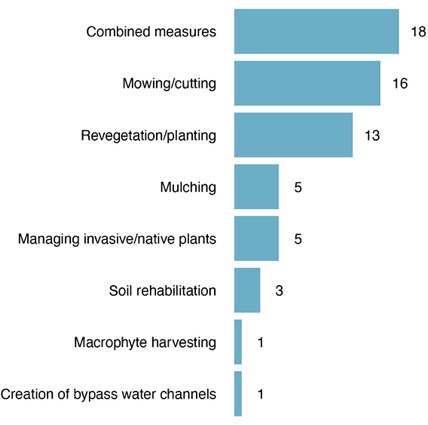

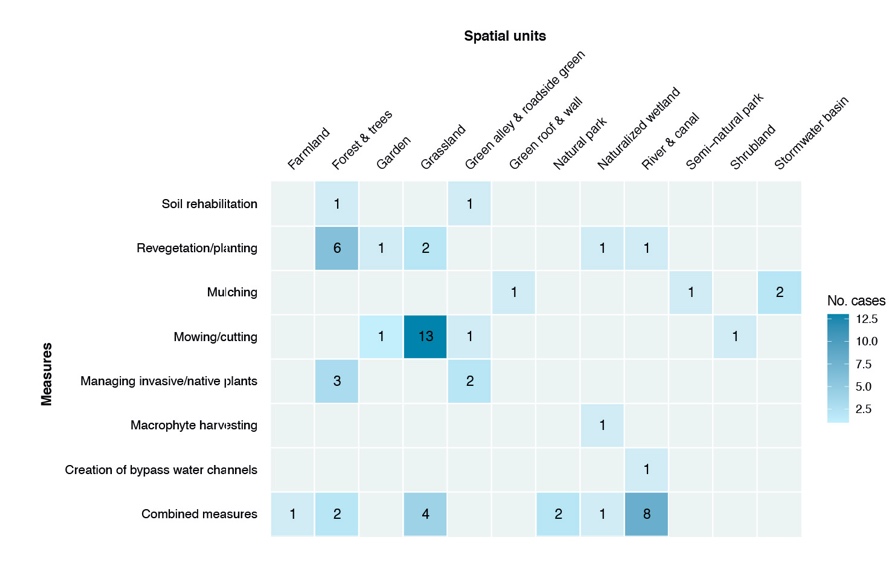

**Fig. S1**. A) Distribution of measure types applied in the units. B) The number of cases per combination of measure type and unit type identified in our database.

1.2 The links between outcome domains and types of nature-based solutions.

Within the vegetation feature, the highest numbers of links were between forest and trees and the outcomes of plant biodiversity (13), microclimate regulation (12), greenhouse gas mitigation (11), and mental health (11). A high number of links were also found linking grassland to plant biodiversity (12). Within the hybrid feature, general parks (17) and semi-natural parks (14) were commonly linked to mental health outcomes, while semi-natural parks were also found to provide microclimate regulation outcomes (11). Within the engineered feature, the highest number of links was between green roofs & walls and microclimate regulation (15). The highest number of links within the water feature were between stormwater basins and flood mitigation (16), as well as water quality enhancement (13).

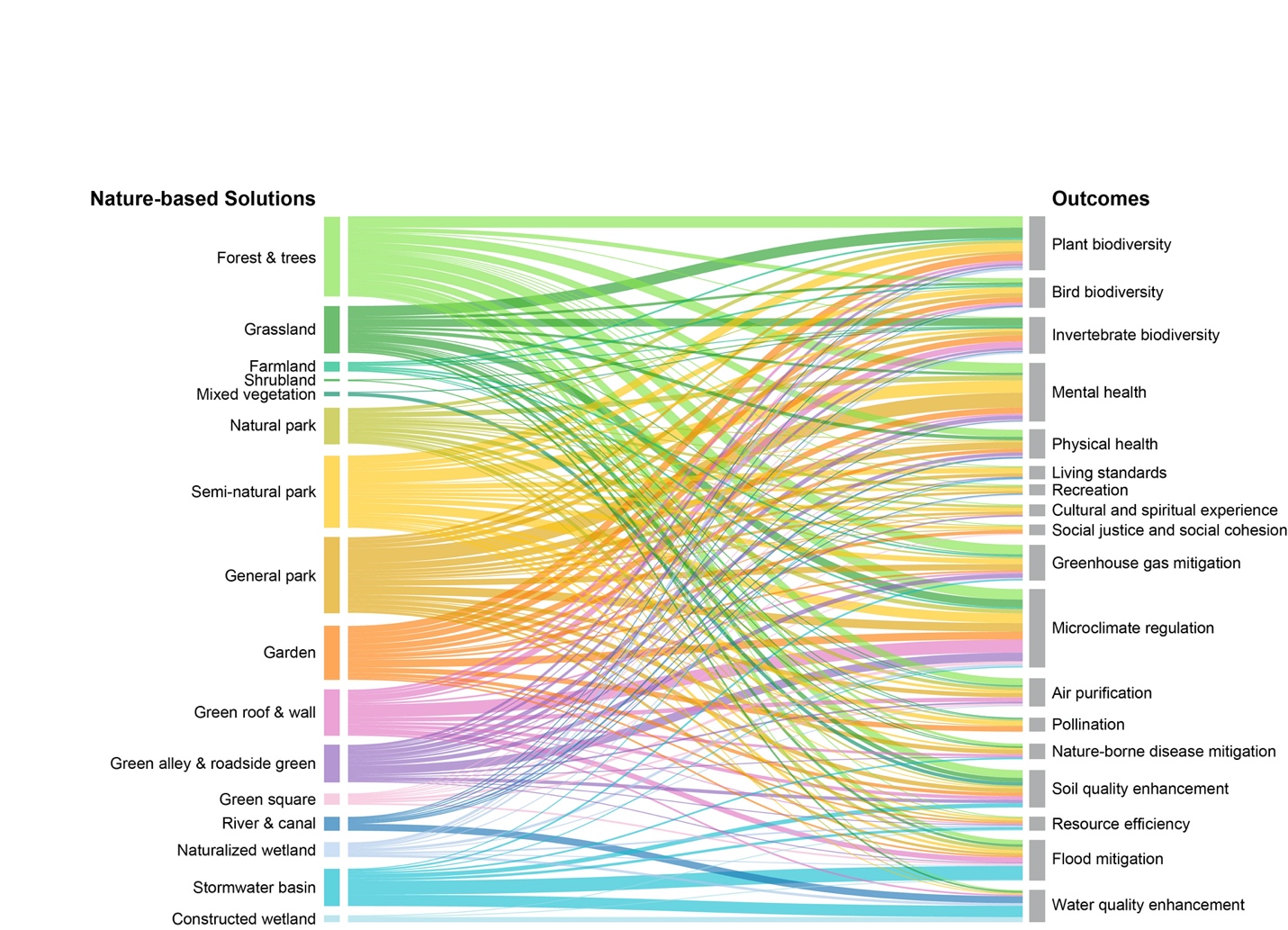

**Fig. S2**. Sankey diagram relating 16 nature-based solutions (NbS) unit types (left) to the 18 most reported outcome domains (right). The data used in the analysis contains 630 outcomes related to 462 NbS types and urban challenges. The thickness of each band corresponds to the number of cases. NbS types were arranged and color-coded according to features: those in green correspond to the vegetation feature, those in yellow to the hybrid feature, those in purple to the engineered feature, and those in blue to the water feature.

### 2 Supplemental figures

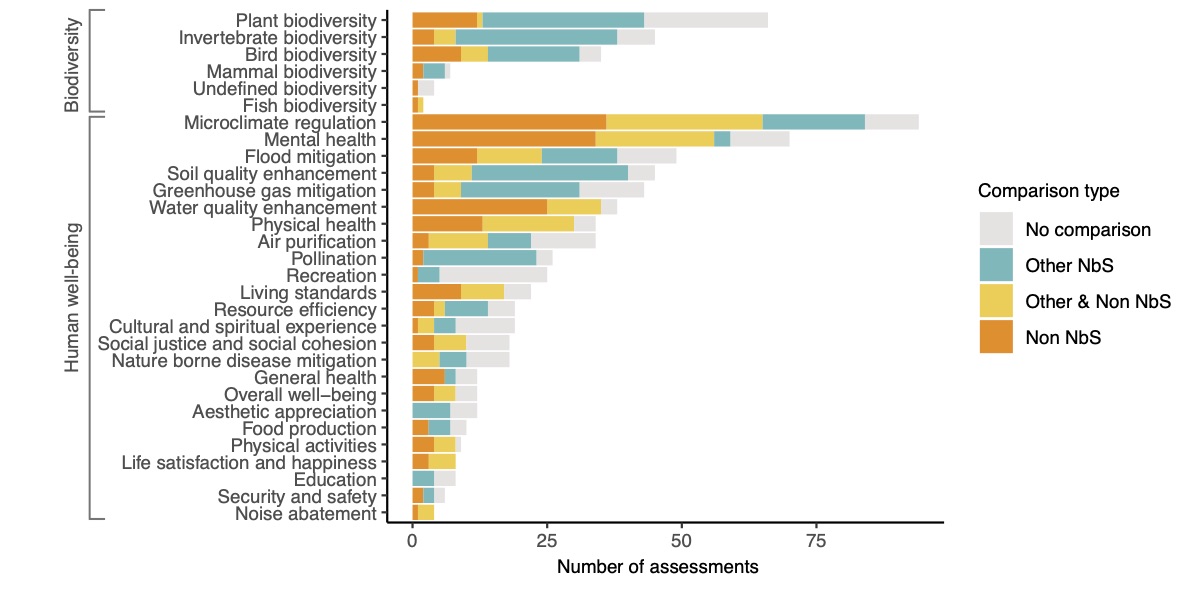

**Fig. S3.** Distribution of types of comparators used across 30 outcome domains.

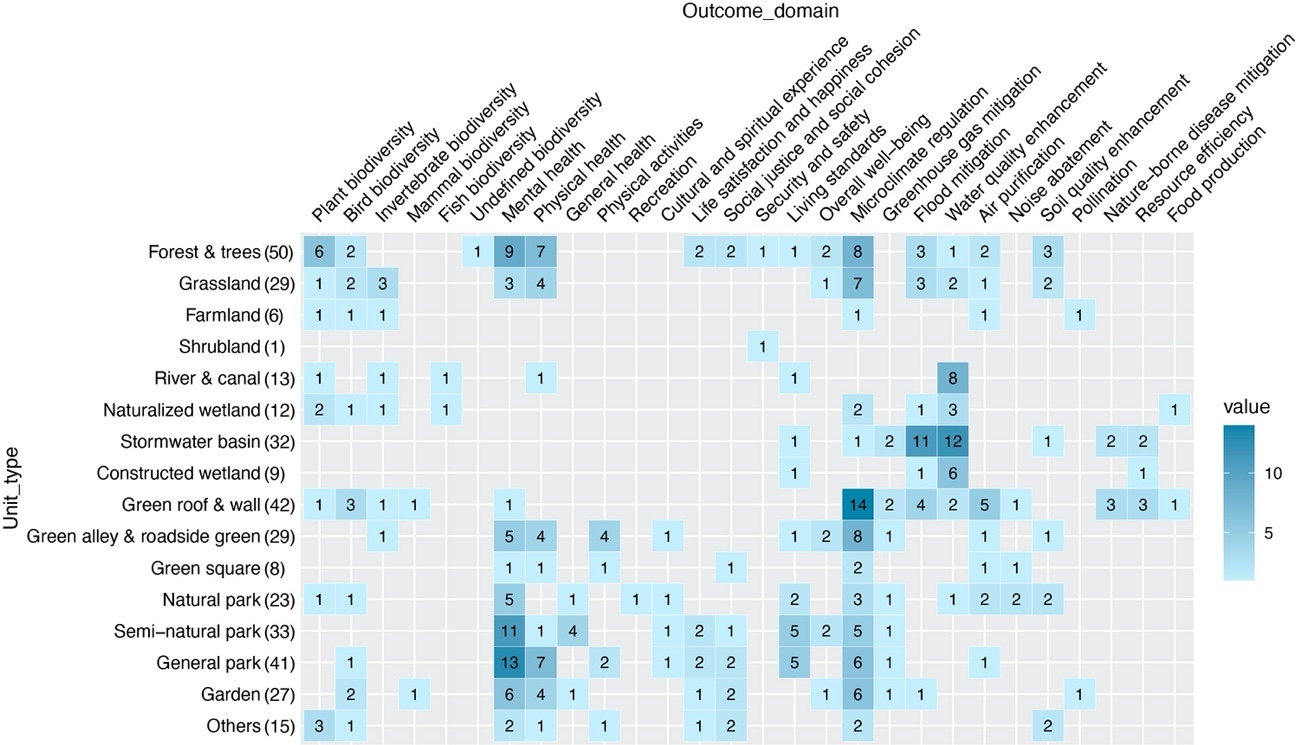

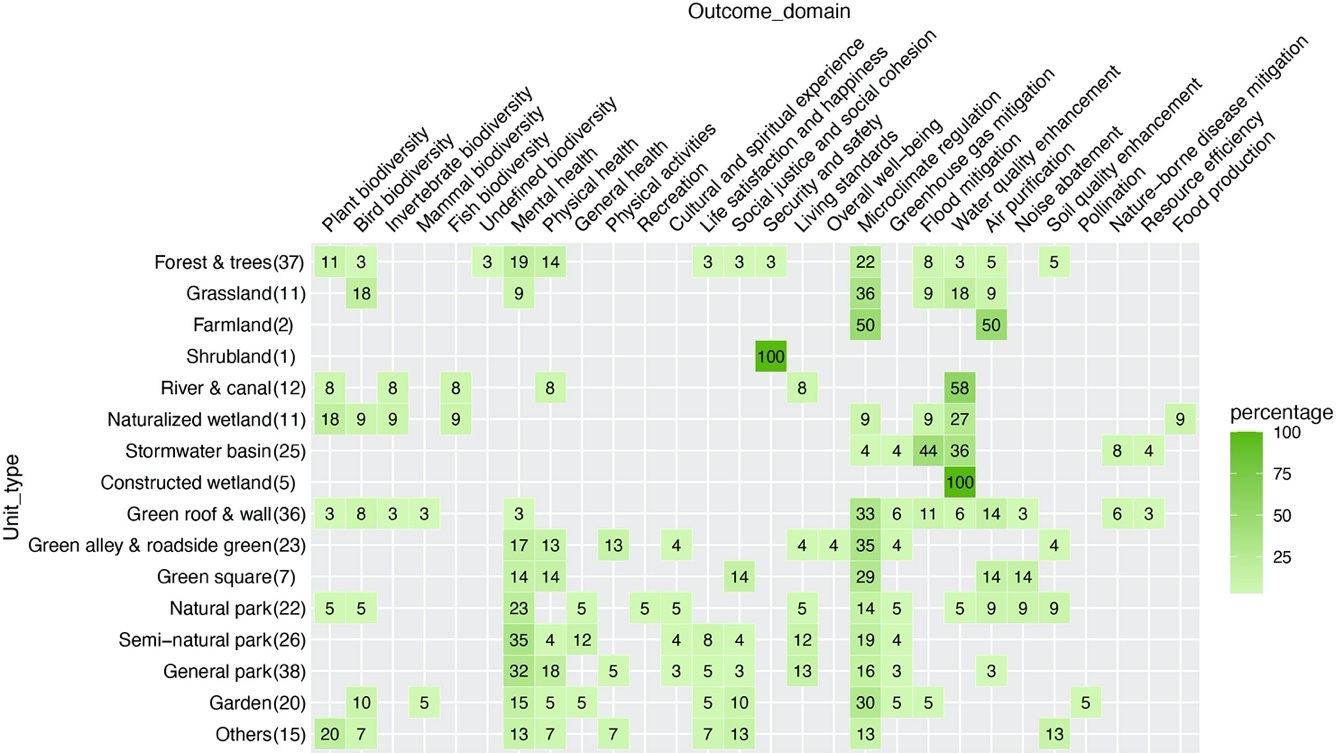

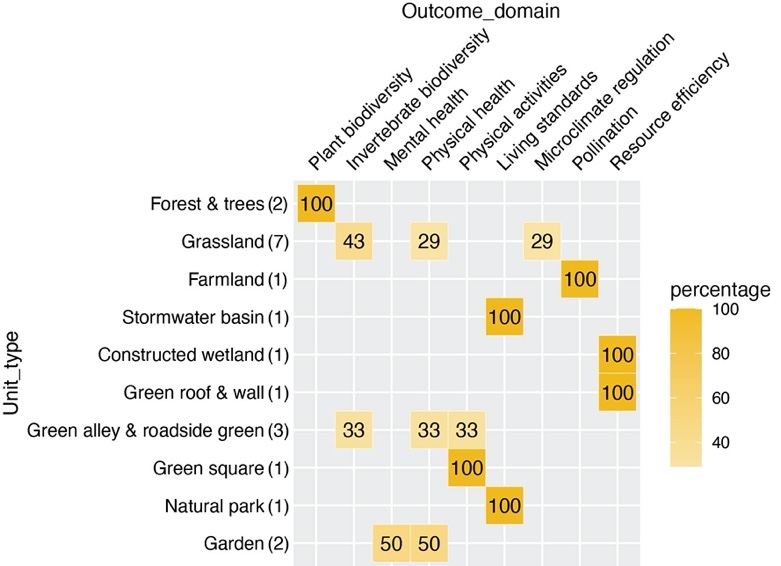

**Fig. S4.** Reported effects per unit type and per outcome domain: A) All directions, shown in number of outcome assessments, B) Positive effects, shown in percentage of reported outcome numbers within each unit type. C) Negative effects, shown in percentage of reported outcome numbers within each unit type.

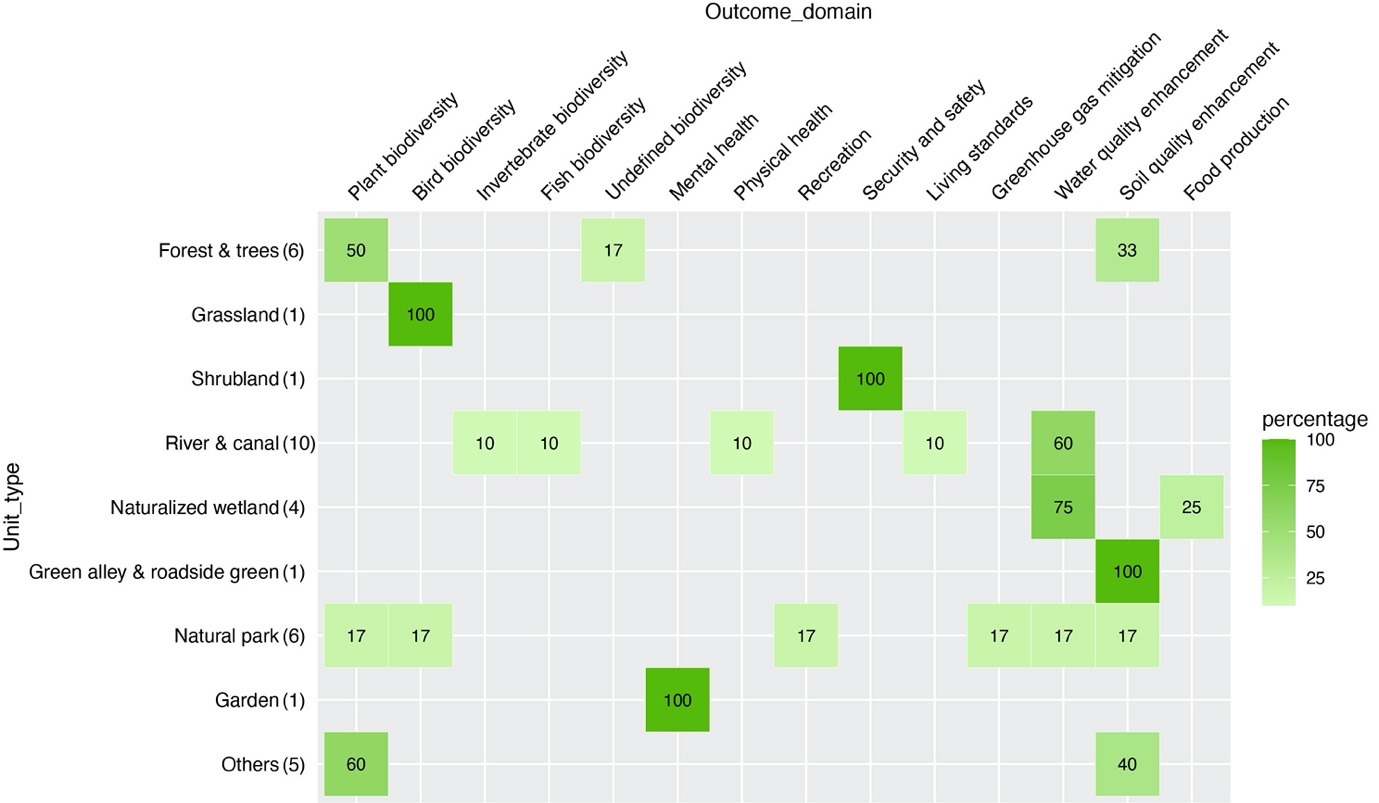

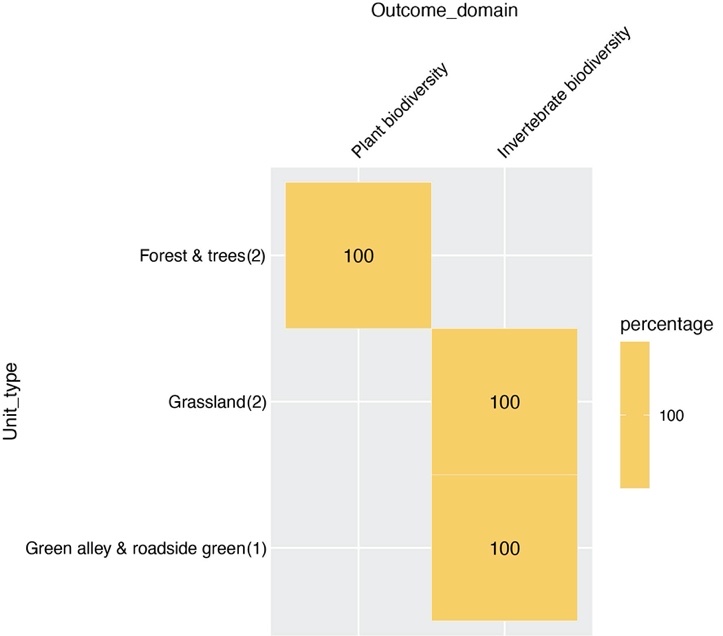

**Fig. S5**. Reported effects per unit type and per outcome domain: A) Positive effects, shown in percentage of reported outcome numbers within each unit type. B) Negative effects, shown in percentage of reported outcome numbers within each unit type.

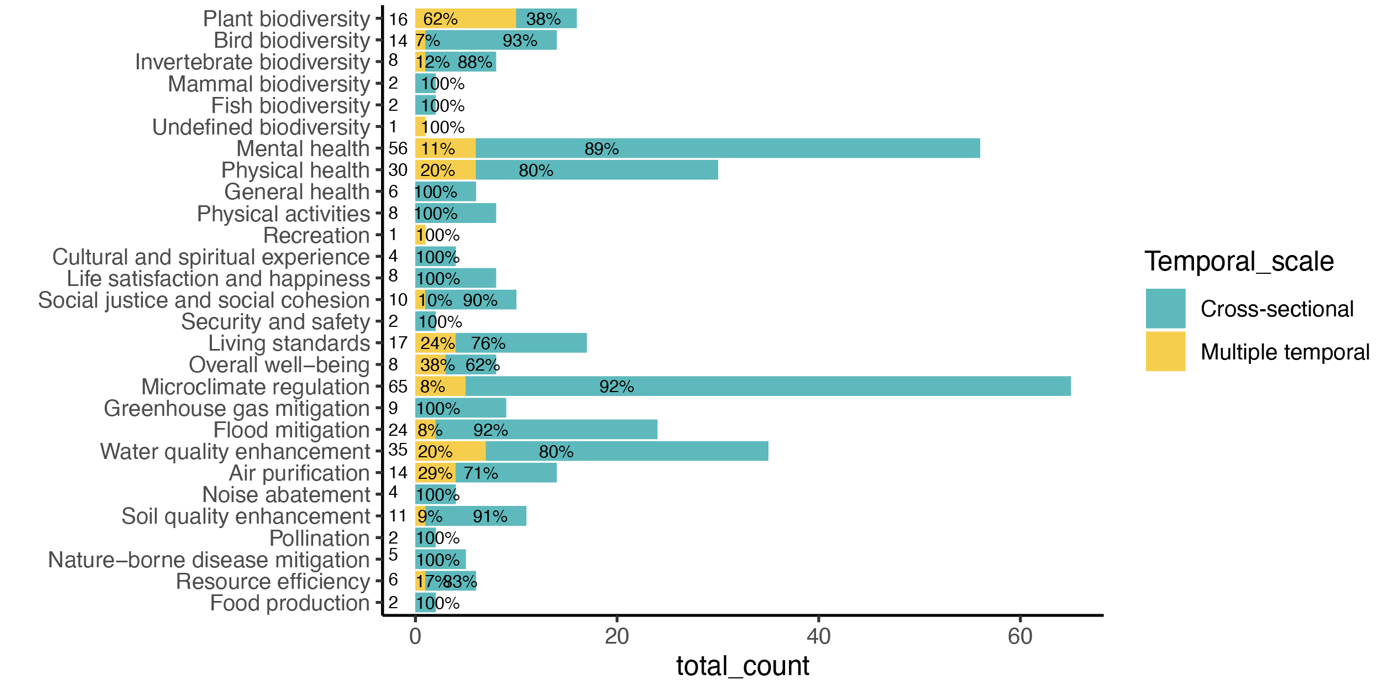

**Fig. S6.** Frequency of cross-sectional effects and multiple effects over time across outcome domains.

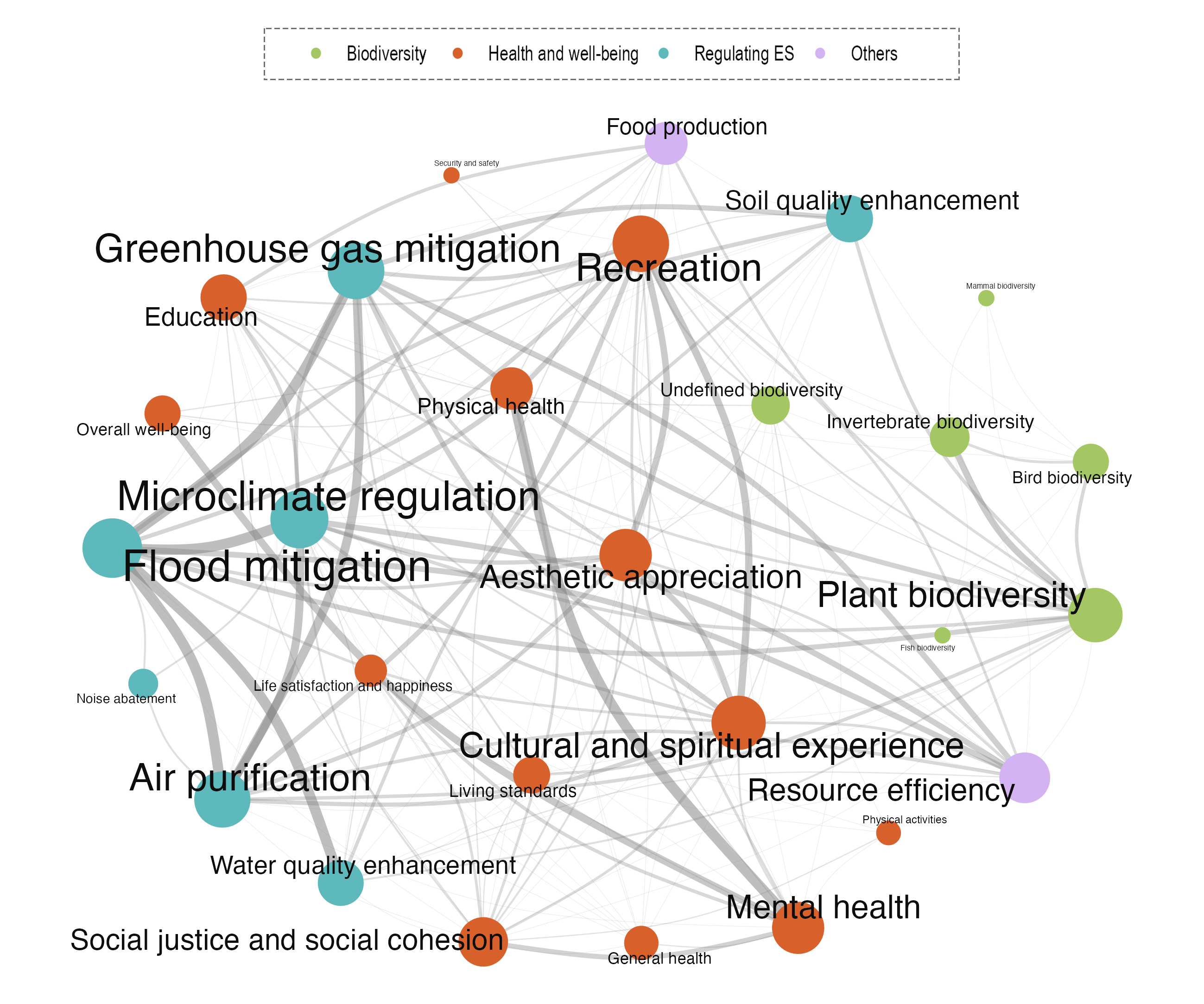

**Fig. S7**. Co-occurrence network of the outcomes. The data includes 387 outcomes generated from 133 cases containing multiple outcomes. The thickness corresponds to the frequencies of the co-occurrence, and the sizes of vertices and texts correspond to the occurrences of outcome domains.

### 3 Supplemental tables

**Table S1.** Geographical distribution of urban nature-based solution cases identified regarding continents, countries, and cities (numbers and % in 547 cases).

**A)** Continent

| Continent | Number | Percentage |
| --- | --- | --- |
| Africa | 16 | 3% |
| Asia | 181 | 33% |
| Europe | 167 | 31% |
| North America | 115 | 21% |
| Oceania | 39 | 7% |
| South America | 28 | 5% |

**B)** The 10 countries with the highest number of cases.

| **Country** | **Number** | **Percentage** |
| --- | --- | --- |
| People's Republic of China | 95 | 17% |
| United States of America | 91 | 17% |
| Australia | 34 | 6% |
| Germany | 30 | 5% |
| United Kingdom | 25 | 5% |
| Brazil | 21 | 4% |
| Spain | 21 | 4% |
| Canada | 20 | 4% |
| India | 15 | 3% |
| Poland | 15 | 3% |

Note: Analysis is based on 546 records by removing one case with multiple countries. The percentage is counted in 547 interventions.

**C) The 10 cities with the highest number of cases.**

| **Cities** | **Number** | **Percentage** |
| --- | --- | --- |
| Beijing | 27 | 5% |
| New York | 22 | 4% |
| Singapore | 13 | 2% |
| Guangzhou | 11 | 2% |
| Toronto | 11 | 2% |
| Adelaide | 11 | 2% |
| Chicago | 9 | 2% |
| Hong Kong | 9 | 2% |
| Melbourne | 8 | 1% |
| Shanghai | 8 | 1% |

Note that 18 cases had multiple locations (multiple) and 21 without mentioning specific locations (NA) not shown in the table.

**Table S2.** Distribution of NbS units in identified urban NbS cases, categorized by features (A) and types (B) (number and % in 547 cases).

**A) Features**

| **Unit feature** | **Number** | **Percentage** |
| --- | --- | --- |
| Hybrid | 225 | 41% |
| Vegetation | 156 | 29% |
| Engineered | 94 | 17% |
| Water | 61 | 11% |
| Others | 11 | 2% |
| Grand Total | 547 | 100% |

**B) Types**

| **Unit types** | **Number** | **Percentage** |
| --- | --- | --- |
| Forest & trees | 86 | 16% |
| Semi-natural park | 72 | 13% |
| General park | 69 | 13% |
| Garden | 58 | 11% |
| Grassland | 52 | 10% |
| Green roof & wall | 49 | 9% |
| Green alley & roadside green | 37 | 7% |
| Stormwater basin | 29 | 5% |
| Natural park | 26 | 5% |
| River & canal | 14 | 3% |
| Naturalized wetland | 11 | 2% |
| Others | 11 | 2% |
| Farmland | 10 | 2% |
| Green square | 8 | 1% |
| Constructed wetland | 7 | 1% |
| Mixed vegetation | 5 | 1% |
| Shrubland | 3 | 1% |
| Grand Total | 547 | 100% |

**Table S3.** Distribution of NbS measures in identified urban NbS cases combining specific measures in units, categorized by features (A) and types (B) (number and % in 62 cases).

**A) Features**

| **Measure features** | **Number** | **Percentage** |
| --- | --- | --- |
| Plant | 35 | 56% |
| Combined measures | 18 | 29% |
| Soil | 8 | 13% |
| Abiotic | 1 | 2% |
| Grand Total | 62 | 100% |

**B) Types**

| **Meature type** | **Number** | **Percentage** |
| --- | --- | --- |
| Combined measures | 18 | 29% |
| Mowing/cutting | 16 | 26% |
| Revegetation/planting | 13 | 21% |
| Mulching | 5 | 8% |
| Managing invasive/native plants | 5 | 8% |
| Soil rehabilitation | 3 | 5% |
| Creation of bypass water channels | 1 | 2% |
| Macrophyte harvesting | 1 | 2% |
| Grand Total | 62 | 100% |

**Table S4.** Distribution of categories of urban challenges in identified urban NbS cases (number and % in 547 cases).

| **Categories** | **Number** | **Percentage** |
| --- | --- | --- |
| Health and well-being | 140 | 26% |
| Biodiversity loss | 103 | 19% |
| Urban heat island | 68 | 12% |
| No specific challenges | 40 | 7% |
| Habitat degradation | 33 | 6% |
| Multiple | 25 | 5% |
| Water pollution and shortage | 20 | 4% |
| Greenhouse gas emission | 19 | 3% |
| Flood risk | 18 | 3% |
| Soil degradation | 17 | 3% |
| Air pollution | 16 | 3% |
| Lack of open green spaces | 13 | 2% |
| Pollinator and pollination loss | 10 | 2% |
| Food insecurity | 5 | 1% |
| Urban shrink and decline | 4 | 1% |
| Public perception and participation | 3 | 1% |
| Urban development issue | 3 | 1% |
| Ecosystem disservices | 3 | 1% |
| Social justice and cohesion issue | 2 | 0% |
| Natural hazards | 2 | 0% |
| Noise pollution | 1 | 0% |
| Cultural heritage protection | 1 | 0% |
| Climate change | 1 | 0% |
| Grand Total | 547 |  |

**Table S5.** Distribution of categories of outcome domains in identified outcomes assessments (number and % in 799 outcomes).

**a) Three broad outcome domains**

| **Broad domains** | **Number** | **Percentage** |
| --- | --- | --- |
| Regulating ecosystem services | 353 | 44% |
| Health and well-being | 255 | 32% |
| Biodiversity | 162 | 20% |
| Others | 29 | 4% |
| Total of the outcomes | 799 | 100% |

**b) 30 outcome domains**

| **Domains** | **Number** | **Percentage** |
| --- | --- | --- |
| Microclimate regulation | 94 | 12% |
| Mental health | 70 | 9% |
| Plant biodiversity | 69 | 9% |
| Flood mitigation | 49 | 6% |
| Soil quality enhancement | 47 | 6% |
| Invertebrate biodiversity | 45 | 6% |
| Greenhouse gas mitigation | 43 | 5% |
| Water quality enhancement | 38 | 5% |
| Bird biodiversity | 35 | 4% |
| Air purification | 34 | 4% |
| Physical health | 34 | 4% |
| Pollination | 26 | 3% |
| Recreation | 25 | 3% |
| Living standards | 22 | 3% |
| Cultural and spiritual experience | 19 | 2% |
| Resource efficiency | 19 | 2% |
| Social justice and social cohesion | 18 | 2% |
| Nature-borne disease mitigation | 18 | 2% |
| Overall well-being | 12 | 2% |
| General health | 12 | 2% |
| Aesthetic appreciation | 12 | 2% |
| Food production | 10 | 1% |
| Physical activities | 9 | 1% |
| Education | 8 | 1% |
| Life satisfaction and happiness | 8 | 1% |
| Mammal biodiversity | 7 | 1% |
| Security and safety | 6 | 1% |
| Noise abatement | 4 | 1% |
| Undefined biodiversity | 4 | 1% |
| Fish biodiversity | 2 | 0% |
| Total of the outcomes | 799 | 100% |

**Table S6.** The number of broad categories of outcomes per NbS unit types (in 630 outcomes included in the linkage analysis).

| **NbS Unit featured groups** | **Unit Types** | **BIO** | **HWB** | **RES** | **Others** | **Grand Total** |
| --- | --- | --- | --- | --- | --- | --- |
| Blue | Constructed wetland | 0 | 0 | 7 | 1 | 8 |
| Blue | Naturalized wetland | 6 | 1 | 8 | 2 | 17 |
| Blue | River & canal | 3 | 5 | 8 | 0 | 16 |
| Blue | Stormwater basin | 0 | 1 | 39 | 3 | 43 |
| Green | Farmland | 5 | 0 | 6 | 0 | 11 |
| Green | Forest & trees | 19 | 23 | 49 | 1 | 92 |
| Green | Grassland | 24 | 7 | 23 | 0 | 54 |
| Green | Mixed vegetation | 1 | 0 | 4 | 0 | 5 |
| Green | Shrubland | 1 | 0 | 1 | 0 | 2 |
| Grey | Green alley & roadside green | 7 | 12 | 23 | 1 | 43 |
| Grey | Green roof & wall | 13 | 2 | 37 | 1 | 53 |
| Grey | Green square | 2 | 3 | 8 | 0 | 13 |
| Hybrid | Garden | 21 | 16 | 23 | 2 | 62 |
| Hybrid | General park | 13 | 36 | 36 | 2 | 87 |
| Hybrid | Natural park | 5 | 15 | 20 | 1 | 41 |
| Hybrid | Semi-natural park | 18 | 31 | 32 | 2 | 83 |
|  | **Total** | **138** | **152** | **324** | **16** | **630** |

**Table S7.** Distribution of types of the comparator used in the outcome assessments (number and % of the 799 outcomes).

| **Type** | **Number of outcomes** | **Percentage** |
| --- | --- | --- |
| Both | 166 | 21% |
| NA | 189 | 24% |
| Non | 207 | 26% |
| Other | 237 | 30% |
| Total | 799 | 100% |

**Table S8.** Distribution of categories of reported effects by Non-NbS comparator.

| Effects | Number of outcomes | Percentage in 799 outcomes | Percentage of 370 reported effects |
| --- | --- | --- | --- |
| Positive | 291 | 36% | 79% |
| Negative | 30 | 4% | 5% |
| Neutral | 29 | 4% | 8% |
| Mixed | 20 | 3% | 8% |
| NA | 429 | 54% | - |
| Total | 799 | 100% |  |

**Table S9.** Distribution of numbers of outcomes within a case ( in number and % of 133 cases reporting multiple outcomes).

| **Number of outcomes** | **Counts of cases** | **Percentage** |
| --- | --- | --- |
| 2 | 78 | 58,6% |
| 3 | 22 | 16,5% |
| 4 | 21 | 15,8% |
| 5 | 4 | 3,0% |
| 6 | 3 | 2,3% |
| 8 | 4 | 3,0% |
| 11 | 1 | 0,8% |
| Total | 133 | 100,0% |

Table S10. Distribution of reported multiple effects within a case (in number and % of 133 cases reporting multiple outcomes). Summarized based on the coding database, by filtering the cases with multiple records in column ‘Performance_non_NbS’ within the same study ID.

| **Number of reported effects within a case** | **Subcategory** | **Count of cases** |
| --- | --- | --- |
| ***Multiple effects*** |  | ***63*** |
|  | 2 and more positive effects (synergy) | 20 |
|  | At least 1 positive and 1 negative effect (trade-off) | 1 |
|  | Other conditions | 42 |
| ***Less than 2 effects*** |  | ***70*** |

Table S11. List of 20 cases with multiple positive effects (summarized based on data used in Table 10 by filtering the cases with 2 and more ‘positive’ records within the column ‘Performance_non_NbS’).

| **Study_ID** | **Outcome_domain** | **Unit_type** | **Urban_challenge** |
| --- | --- | --- | --- |
| S0020 | Plant biodiversity | Forest & trees | Habitat degradation |
| S0020 | Undefined biodiversity | Forest & trees | Habitat degradation |
| S0048 | Mental health | General park | Health and well-being |
| S0048 | Life satisfaction and happiness | General park | Health and well-being |
| S0077 | Invertebrate biodiversity | River & canal | Multiple |
| S0077 | Water quality enhancement | River & canal | Multiple |
| S0101 | Flood mitigation | Green roof & wall | Flood risk |
| S0101 | Food production | Green roof & wall | Flood risk |
| S0113 | Physical health | General park | Health and well-being |
| S0113 | Mental health | General park | Health and well-being |
| S0123 | Physical health | Forest & trees | Health and well-being |
| S0123 | Mental health | Forest & trees | Health and well-being |
| S0218 | Physical activities | General park | Health and well-being |
| S0218 | Physical health | General park | Health and well-being |
| S0242 | Water quality enhancement | Stormwater basin | Multiple |
| S0242 | Flood mitigation | Stormwater basin | Multiple |
| S0312 | Life satisfaction and happiness | Semi-natural park | Health and well-being |
| S0312 | General health | Semi-natural park | Health and well-being |
| S0312 | Mental health | Semi-natural park | Health and well-being |
| S0316 | Food production | Naturalized wetland | Food insecurity |
| S0316 | Water quality enhancement | Naturalized wetland | Food insecurity |
| S0323 | Resource efficiency | Stormwater basin | Flood risk |
| S0323 | Greenhouse gas mitigation | Stormwater basin | Flood risk |
| S0350 | Mental health | Natural park | Health and well-being |
| S0350 | Cultural and spiritual experience | Natural park | Health and well-being |
| S0382 | Plant biodiversity | Natural park | Habitat degradation |
| S0382 | Water quality enhancement | Natural park | Habitat degradation |
| S0382 | Bird biodiversity | Natural park | Habitat degradation |
| S0382 | Soil quality enhancement | Natural park | Habitat degradation |
| S0382 | Greenhouse gas mitigation | Natural park | Habitat degradation |
| S0382 | Recreation | Natural park | Habitat degradation |
| S0390 | Physical activities | Others | Health and well-being |
| S0390 | Mental health | Others | Health and well-being |
| S0390 | Life satisfaction and happiness | Others | Health and well-being |
| S0390 | Social justice and social cohesion | Others | Health and well-being |
| S2014 | Mental health | Green alley & roadside green | Health and well-being |
| S2014 | Microclimate regulation | Green alley & roadside green | Health and well-being |
| S2024 | Flood mitigation | Stormwater basin | Water pollution and shortage |
| S2024 | Water quality enhancement | Stormwater basin | Water pollution and shortage |
| S2047 | Bird biodiversity | Green roof & wall | Biodiversity loss |
| S2114 | Plant biodiversity | Others | Urban shrink and decline |
| S2114 | Soil quality enhancement | Others | Urban shrink and decline |
| S2047 | Invertebrate biodiversity | Green roof & wall | Biodiversity loss |
| S2120 | General health | Garden | Health and well-being |
| S2120 | Physical health | Garden | Health and well-being |
| S2120 | Mental health | Garden | Health and well-being |
| S2120 | Social justice and social cohesion | Garden | Health and well-being |
| S2162 | Physical health | General park | Health and well-being |
| S2162 | Mental health | General park | Health and well-being |
